# SLC19A1 is a cyclic dinucleotide transporter

**DOI:** 10.1101/539767

**Authors:** Rutger D. Luteijn, Shivam A. Zaver, Benjamin G. Gowen, Stacia Wyman, Nick Garelis, Liberty Onia, Sarah M. McWhirter, George E. Katibah, Jacob E. Corn, Joshua J. Woodward, David H. Raulet

## Abstract

The accumulation of DNA in the cytosol serves as a key immunostimulatory signal associated with infections, cancer and genomic damage^1,2^. Cytosolic DNA triggers immune responses by activating the cGAS/STING pathway^3^. The binding of DNA to the cytosolic enzyme cGAMP synthase (cGAS), activates its enzymatic activity, leading to the synthesis of a second messenger, cyclic[G(2’,5’)pA(3’,5’)] (2’3’-cGAMP)^4–8^. 2’3’-cGAMP, a cyclic dinucleotide (CDN), activates the protein ‘stimulator of interferon genes’ (STING)^9^, which in turn activates the transcription factors IRF3 and NF-κB promoting the transcription of genes encoding type I interferons and other cytokines and mediators that stimulate a broader immune response. Exogenous 2’3’-cGAMP and other CDNs, including CDNs produced by bacteria and synthetic CDNs used in cancer immunotherapy, must traverse the cell membrane to activate STING in target cells. How these charged CDNs pass through the lipid bilayer is unknown. Here we used a genome-wide CRISPR interference screen to identify the reduced folate carrier SLC19A1 as the major CDN transporter for uptake of synthetic and naturally occurring CDNs. CDN uptake and functional responses are inhibited by depleting SLC19A1 from cells and enhanced by overexpressing *SLC19A1*. In both cell lines and primary cells *ex vivo*, CDN uptake is inhibited competitively by folate and blocked by the SLC19A1 inhibitor sulfasalazine, a medication approved for the treatment of inflammatory diseases. The identification of SLC19A1 as the major transporter of CDNs into cells has far reaching implications for the immunotherapeutic treatment of cancer^10^, transport of 2’3’-cGAMP from tumor cells to other immune cells to trigger the anti-tumor immune response^11^, host responsiveness to CDN-producing pathogenic microorganisms^12^, and potentially in certain inflammatory diseases.

## Main text

The cGAS/STING pathway senses cytosolic DNA originating from viruses and bacteria^9^ as well as CDNs produced by certain bacteria^13–15^. Notably, the STING pathway is also activated by cytosolic self DNA, which accumulates in cells in certain autoinflammatory disorders, including Aicardi–Goutieres Syndrome and systemic lupus erythematosus^16–19^. Furthermore, cytosolic DNA accumulates in cells subjected to DNA damage, as occurs in tumor cells, resulting in activation of the cGAS/STING pathway and the initiation of an anti-tumor immune response^20^. Recently, we revealed that 2’3’-cGAMP can be transferred from tumor cells to immune cells *in vivo*, prompting the activation of the immune response^11^. Furthermore, synthetic STING agonists, such as 2’3’-RR CDA, an analogue of 2’3’-cGAMP (Fig. S1)^21^, can greatly enhance the anti-tumor immune response when delivered directly into the tumor microenvironment in mouse models of cancer, causing tumor regressions^10,22^. 2’3’-RR CDA and other synthetic CDNs are currently being tested in clinical trials as cancer immunotherapies. However, a critical outstanding question is the mechanism of transport of CDNs into cells of the immune system^23^. CDNs may be incorporated into cells via gap junctions, membrane fusions, or by incorporation into viral particles^24–27,28^. but none of these mechanisms explain (systemic) immune activation by extracellular CDNs. To systematically identify the genes involved in cytosolic transport of CDNs, we performed a genome-wide CRISPR interference screen in the monocytic THP-1 cell line.

To visualize STING activation, THP-1 cells were transduced with a CDN-inducible reporter construct (Fig. 1a). The reporter was composed of Interferon Stimulatory Response Elements (ISRE) and a mouse minimal IFN-β promoter that drives the expression of tdTomato upon hIFN-β or CDN exposure (Fig. 1b). In line with previous results, the synthetic CDN 2’3’-RR CDA induced a more potent response than 2’3’-cGAMP^10^ even when applied at a lower concentration. The response to both CDNs was several fold higher than the response to hIFN-β, and was completely dependent on STING expression (Fig. 1b), implying that the reporter primarily reported cell-intrinsic STING activity. To interrogate the approximately 20,000 human genes for their role in CDN-induced reporter expression, we performed a genome-wide CRISPR interference (CRISPRi) forward genetic screen in THP-1 cells. We generated a stable line of THP-1 cells expressing dCas9-BFP-KRAB, which was validated and expanded before transducing the cells with the CRISPRi v2 library at a low multiplicity of infection (see Methods). The CRISPRi library of cells was stimulated either with 2’3’-RR CDA, or with 2’3’-cGAMP, using concentrations that resulted in 90% reporter-positive cells. The highest expressing 25% and lowest expressing 25% of stimulated cells in each library were sorted by flow cytometry, DNA isolated, and gRNA sequences from each population, and unsorted cells, were amplified and DNA from each of these populations as well as from unsorted cells was deep-sequenced to identify the targeted genes in each population (Fig. 1c and Fig. S2). The fold enrichment and depletion of gRNAs in the hypo-responsive population versus the hyper-responsive population was calculated for each screen (Fig. S3, Table S1 and Table S2). We also integrated multiple gRNAs per gene using Mageck (see Methods) comparing the hyporesponsive and hyperresponsive populations calculated as robust rank aggregations scores and depicted in Fig. 1d and e. Similar results were obtained when each sorted population was compared to unsorted cells (Table S2). The two screens yielded many common hits, but there were some differences, such as numerous hits in the 2’3’-cGAMP screen including STAT2, IRF9, IFNAR1, and IFNAR2 (Table S2). Hence, the 2’3’-RR CDA screen may have been mostly dependent on intrinsic STING signaling, whereas the 2’3’-cGAMP screen may have been partly dependent on autocrine/paracrine IFN-β signaling.

**Figure 1.**
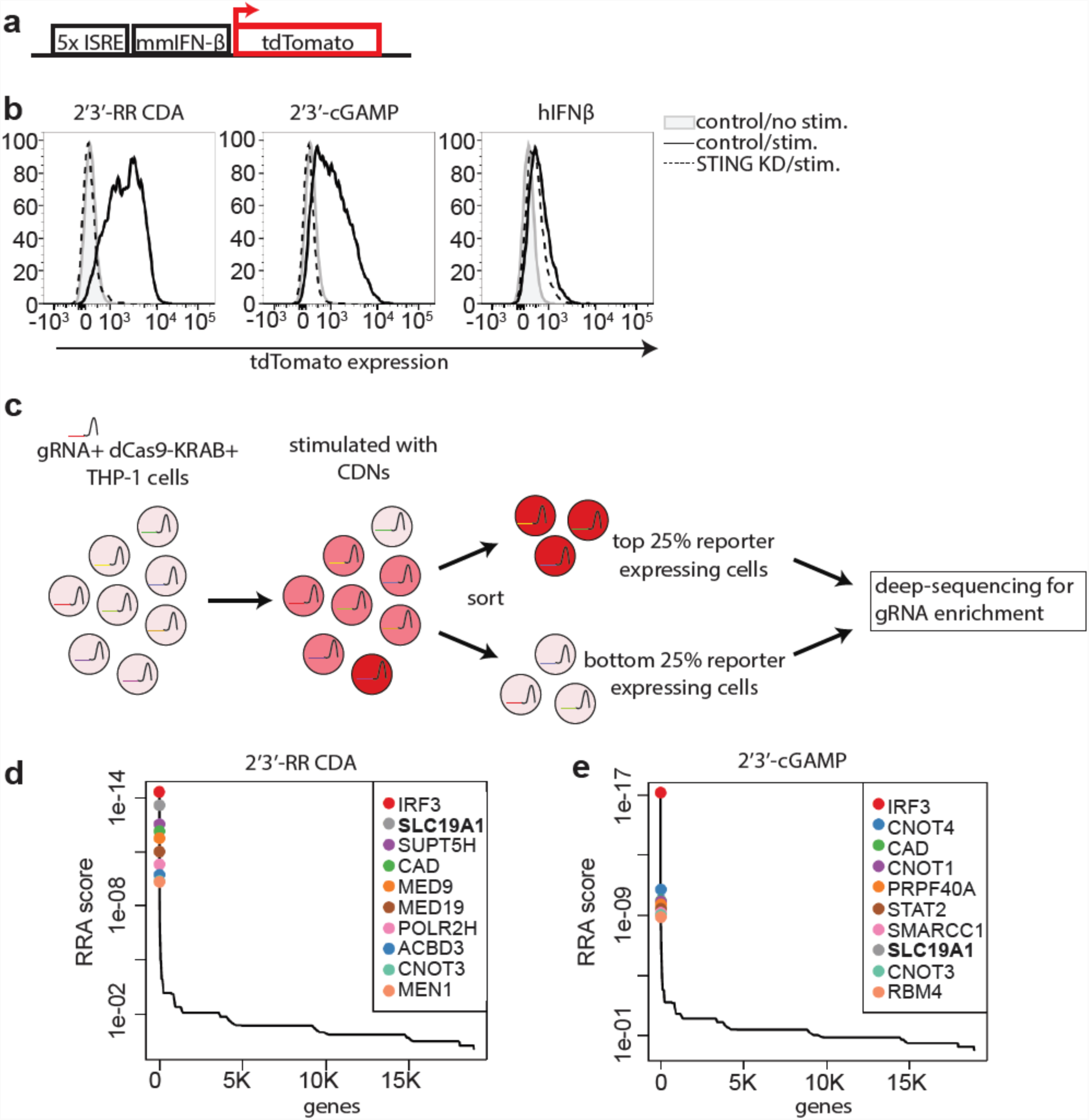
Genome-wide CRISPRi screen for host factors necessary for cyclic dinucleotide (CDN) stimulation. **a,** schematic overview of tdTomato-reporter. tdTomato expression is driven by interferon-stimulatory response elements (ISRE) followed by a mouse minimal interferon beta (mmIFN-β) promoter. **b,** Control THP-1 cells and *STING*-depleted THP-1 cells were incubated with 2’3’-RR CDA (1.67 µg/ml), 2’3’-cGAMP (10 µg/ml) or hIFN-β (100 ng/ml). After 20h, tdTomato reporter expression was analyzed by flow cytometry. Data are representative of three independent experiments with similar results. **c,** Schematic overview of the genome-wide CRISPRi screen. A genome-wide library of CRISPRi guide RNA (gRNA)-expressing THP-1 cells was stimulated with CDNs. 20h after stimulation, cells were sorted into a tdTomato-low group (lowest 25% of cells) and a tdTomato-high group (highest 25% of cells). DNA from the sorted cells was deep sequenced to reveal gRNA enrichment in the two groups. **d-e**, Distribution of the robust rank aggregation (RRA) score in the comparison of hits enriched in the reporter-low versus reporter-high groups of THP-1 cells stimulated with (d) 2’3’-RR CDA or (e) 2’3’-cGAMP. Each panel represents combined results of two independent screens.

In both CDN screens, the top hits in the hypo-responsive population (i.e. the genes most important for robust responses to CDNs) included the transcription factor IRF3, which acts directly downstream of STING. One of the five gRNAs for STING itself was also enriched in hyporesponsive cells from both screens, though the other STING gRNAs were not, presumably because they were ineffective at interfering with STING expression (Table S1). Other significant hits included genes involved in transcription, splicing, and immune modulation (Table S2).

One of the most significant hits in both screens was the *SLC19A1* gene. SLC19A1 is a cell surface transporter known as the reduced folate carrier. SLC19A1 and another transporter, SLC46A1, are responsible for uptake of folate from the extracellular environment^29^. To validate the role of *SLC19A1* in CDN stimulation, the top two enriched *SLC19A1*-targeting gRNAs from the 2’3’-RR CDA screen were used to stably deplete *SLC19A1* in THP-1 cells expressing dCas9-KRAB (Fig. S4a). *SLC19A1*-depleted cells grew normally and appeared healthy, suggesting that other folate transport mechanisms fully suffice in *SLC19A1*-deficient cells. *SLC19A1*-depleted and, for comparison, *IRF3*-depleted cells (Fig. S4b) were stimulated with 2’3’-cGAMP, 2’3’-RR CDA, cyclic [A(3’,5’)pA(3’,5’)] (3’3’ CDA, a bacterial CDN) or hIFN-β, and reporter induction was measured 20 h later (Fig. 2a). Responses to 2’3’-cGAMP, 2’3’-RR CDA and 3’3’-CDA were each strongly inhibited in *IRF3*- and *SLC19A1*-depleted cells (Fig. 2b), whereas stimulation by hIFN-β was not affected (Fig. 2b). Restoration of *SLC19A1* expression by transduction of a cDNA expression vector rescued CDN responsiveness without affecting stimulation by hIFN-β (Fig. 2c).

**Figure 2.**
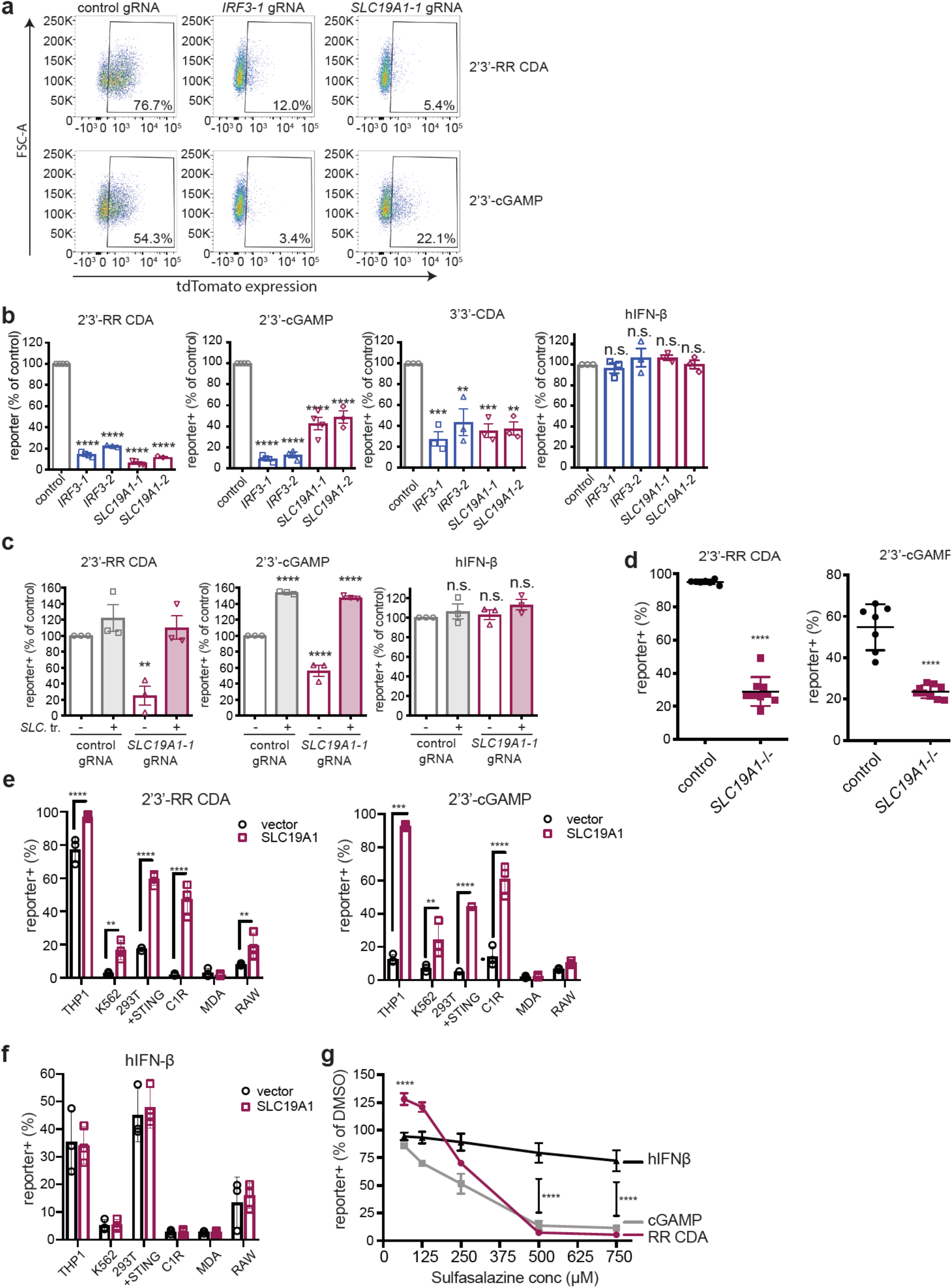
SLC19A1 is required for CDN-induced reporter expression. **a,** dCas9-KRAB-expressing THP-1 cells transduced with non-targeting gRNA (control), *IRF3-1* gRNA or *SLC19A1-1* gRNA were exposed to 2’3’-RR CDA (1.67 µg/ml) or 2’3’-cGAMP (10 µg/ml). 20h later, tdTomato expression was analyzed by flow cytometry. Representative dot plots of three independent experiments are shown. **b,** THP-1 cells expressing the indicated CRISPRi gRNAs or non-targeting gRNA (control), were stimulated with indicated 2’3’-RR CDA (1.67 µg/ml), 2’3’-cGAMP (10 µg/ml), 3’3’ CDA (20 µg/ml) or hIFN-β (100 ng/ml). After 18-22h, tdTomato expression was quantified as in (a). Combined results of three independent experiments are shown. **c,** Control THP-1 cells and *SLC19A1-1* gRNA expressing THP-1 cells transduced with *SLC19A1* (SLC. tr.) were exposed to 2’3’-RR CDA (1.67 µg/ml), 2’3’-cGAMP (15 µg/ml) or hIFN-β (100 ng/ml). After 18-22h, tdTomato reporter expression was quantified. Combined results of three independent experiments are shown. **d,** Control THP-1 cells (7 clonal lines) and *SLC19A1*-/-cells (9 clonal lines) were exposed to 2’3’-RR CDA (2.22 µg/ml), 2’3’-cGAMP (10 µg/ml), and tdTomato reporter expression was analyzed by flow cytometry 20h after stimulation. **e,** Various cell lines expressing a control vector or an *SLC19A1* expression vector were stimulated with 2’3’-RR CDA (1.67 µg/ml) or 2’3’-cGAMP (10 µg/ml). After 20h, reporter expression was quantified by flow cytometry. **f,** Various cell lines expressing a control vector or an *SLC19A1* expression vector were stimulated with hIFN-β (100 ng/ml) or murine IFN-β (100 ng/ml) in the case of RAW cells. After 20h, reporter expression was quantified by flow cytometry. **g,** THP-1 cells were incubated with increasing concentrations of 2’3’-RR CDA, 2’3’-cGAMP or hIFN-β in the presence of the SLC19A1 inhibitor sulfasalazine or DMSO as vehicle control. After 18h, tdTomato reporter expression was analyzed by flow cytometry. For each concentration of sulfasalazine, reporter expression in treated cells was compared to reporter expression in cells treated with the same amount of vehicle (DMSO). In panels b and c, e, f, and g, error bars represent ± SE of three biological replicates. Statistical analysis was performed using one-way ANOVA followed by Tukey’s post-test (b and c), unpaired two-tailed Student’s t tests for (d), two-way ANOVA followed by uncorrected Fisher’s LSD tests (e and f), and two-way ANOVA followed by Tukey’s post-tests to compare the significance between the CDNs and hIFN-β in (g). \**P* ≤ 0.05; \*\**P* ≤ 0.01;\*\*\**P* ≤0.001; \*\*\*\**P* ≤ 0.0001; n.s. not significant

As an alternative approach to corroborate the role of *SLC19A1* in CDN responses, the conventional CRISPR/Cas9 system was used to target a coding exon in order to generate loss of function mutations in the *SLC19A1* gene. Disruption of the *SLC19A1* gene was confirmed by genomic PCR, TA-cloning and sequencing for nine *SLC19A1^-/-^* clones (see Methods). These clones were all significantly less sensitive to CDN stimulation when compared to seven control clones that received a non-targeting gRNA (Fig. 2d).

Importantly, *SLC19A1* overexpression robustly increased CDN responsiveness in THP-1 cells as well as in cell lines that normally responded poorly or not at all to CDN stimulation, including C1R, K562, 293T (pre-transduced with STING), and RAW macrophage cell lines (Fig. 2e and f). Taken together, our data show reduced responses to CDNs in *SLC19A1*-deficient cells and much amplified responses in cells overexpressing *SLC19A1*, as might be expected for a CDN transporter. Together, these data support a central role of the SLC19A1 transporter in responses to several cyclic dinucleotides, including the mammalian CDN 2’3’-cGAMP.

Based on our findings, we tested whether the drug sulfasalazine (SSZ), a non-competitive inhibitor of SLC19A1^30^, would block stimulation by CDNs. THP-1 reporter cells were exposed to various concentrations of SSZ or DMSO vehicle in the presence of 2’3’-cGAMP, 2’3’-RR CDA, or hIFN-β. Responses to both CDNs were robustly inhibited with increasing concentrations of SSZ, whereas responses to hIFN-β stimulation were only modestly inhibited (Fig. 2g). The concentrations required for inhibition were only modestly higher than those that inhibit uptake of folate derivatives in another study^30^. Surprisingly, at lower concentrations, SSZ modestly enhanced stimulation by 2’3’-RR CDA, but had no effect on stimulation by 2’3’ cGAMP (Fig. 2g).

The effect of SLC19A1 on reporter induction by CDNs led us to test the impact of SLC19A1 deficiency on endogenous transcriptional targets downstream of STING, including the genes encoding the chemokines CCL5 and CXCL10, which are direct targets of IRF3^31,32^. In control cells, *CCL5* and *CXCL10* gene expression was highly elevated 5h after 2’3’-RR CDA stimulation. In cells depleted of *IRF3, SLC19A1* or *STING*, chemokine expression was strongly inhibited, indicating that SLC19A1 action is necessary for CDN-induced effects, including those downstream of STING (Fig. 3a and b).

**Figure 3.**
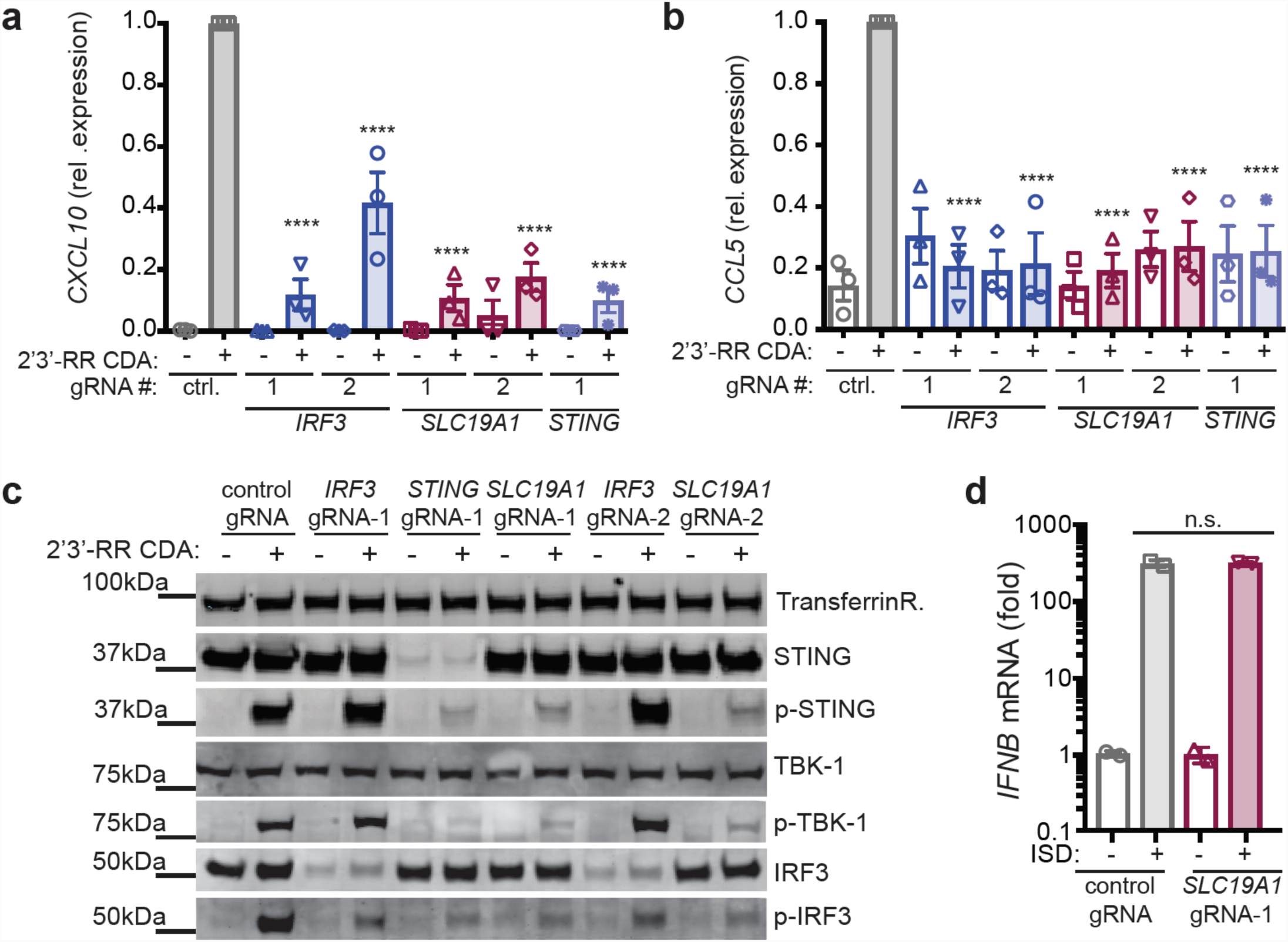
SLC19A1 is critical for STING activation by CDNs. **a, b,** Induction of *CXCL10* (a) or *CCL5* (b) mRNA in control (non-targeting gRNA) THP-1 cells or THP-1 cells expressing the indicated CRISPRi gRNAs after 5h stimulation with 5 µg/ml 2’3’-RR CDA. **c,** Immunoblot analysis of (phospho-) protein expression in control THP-1 cells or THP-1 cells expressing the indicated CRISPRi gRNAs. Cells were stimulated for 2h with 10 µg/ml 2’3’-RR CDA or left unstimulated. TransferrinR.: Transferrin receptor; p-TBK1: TKB1 phosphorylated at position Ser172; p-IRF3: IRF3 phosphorylated at position Ser296; p-STING: STING phosphorylated at position Ser366. Immunoblots are representative of two independent experiments with similar results. **d,** Control THP-1 cells or SLC19A1-depleted THP-1 cells were transfected with 3 εg interferon-stimulatory DNA (ISD) for 3h and the induction of *IFNB* mRNA was measured by RT-qPCR. In panels a, and b: error bars represent ± SE of at least three biological replicates, In panel d: error bar represents ± SE of two biological replicates. Statistical analysis was performed using a one-way ANOVA followed by Dunnett’s post-test for the comparison of the CDN-stimulated *IRF3, SLC19A1*, and *STING*-depleted cell lines to the control CDN-stimulated cells in (a) and an unpaired two-tailed Student’s t test for (d). \*\*\*\**P* ≤ 0.0001; n.s. not significant.

To directly assess the effect of SLC19A1 on STING pathway activation (Fig. S5), we evaluated phosphorylation of STING, IRF3 and TBK1 in control (non-targeting gRNA) versus CRISPRi-depleted cells by immunoblotting (Fig. 3c). Within 2 hours after stimulation with 2’3’-RR CDA, phosphorylation of STING, IRF3, and TBK1 were each significantly elevated in control THP-1 cells. In *IRF3*-depleted cells, phosphorylation of upstream signaling components STING and TBK1 was not affected, whereas in *STING*-depleted cells phosphorylation of both TBK1 and IRF3 was nearly ablated. *SLC19A1*-targeted cells showed major defects in phosphorylation of STING, TBK1 and IRF3, supporting the conclusion that *SLC19A1* acts upstream of STING. Notably, protein levels of STING, TBK1, and IRF3 were unaltered in *SLC19A1*-depleted cells, indicating that SLC19A1 does not influence stability or degradation of STING pathway components.

To further exclude a general defect in STING activation caused by *SLC19A1*-depletion, the cGAS/STING pathway was directly triggered intracellularly by transfecting cells with interferon stimulatory DNA (ISD). ISD transfection of both WT and *SLC19A1*-depleted THP-1 cells resulted in a strong and equal induction of *IFNB* gene expression (Fig. 3d). Thus, STING functioned normally in *SLC19A1*-depleted cells when DNA was introduced directly into the cytosol by transfection.

The finding that *SLC19A1* was not essential when DNA was transfected into cells suggested that SLC19A1 may function by transporting CDNs into cells. Therefore, we enzymatically synthesized [^32^P] 2’3’-cGAMP, which we confirmed by TLC and DRaCALA^33^ binding analysis (Fig. S6a and b). We next monitored 2’3’-cGAMP uptake by cells expressing different levels of SLC19A1. SLC19A1 overexpression greatly enhanced uptake of [^32^P] 2’3’-cGAMP by THP-1 cells (Fig. 4a) and C1R cells (Fig. S7a). Conversely, *SLC19A1*-depletion reduced the uptake of ^32^P 2’3’-cGAMP in THP-1 cells (Fig. 4a). We next sought to determine the specificity for 2’3’-cGAMP uptake in THP-1 cells. Addition of excess, unlabeled bacterial-derived 3’3’-linked cyclic di-nucleotides as well as host-derived 2’3’-cGAMP to cell culture media completely inhibited [^32^P] 2’3’-cGAMP uptake by THP-1 cells, suggesting that cyclic di-nucleotide interactions with the transporter are not highly specific for the 2’3’ linkage or the specific nucleotides (Fig. 4b). In quantitative competition ligand uptake assays, unlabeled 2’3’-cGAMP inhibited uptake of [^32^P] 2’3’-cGAMP with an IC_50_ of 1.89 ± 0.11 µM, in line with the reported affinity of SLC19A1 for methotrexate and other folates (Fig. 4c)^34^. As SLC19A1 was first described as a folate transporter, we performed similar competition experiments using excess, unlabeled folic acid, and we also tested an inhibitor of folate uptake by SLC19A1, sulfasalazine. Remarkably, both folic acid and sulfasalazine inhibited [^32^P] 2’3’-cGAMP uptake with IC_50_’s of 4.79 ± 0.08 µM and 2.06 ± 0.17 µM, respectively (Fig. 4d and e). We extended the study by asking whether folic acid or sulfasalazine inhibited uptake of [^32^P] 2’3’-cGAMP in other cell types. We found that the addition of excess folate and sulfasalazine to cell cultures abrogated [^32^P] 2’3’-cGAMP uptake by U937 monocytes as well as primary murine peritoneal leukocytes and splenocytes (Fig. 4f, S7b). Taken together these results suggested that uptake of the mammalian CDN 2’3’-cGAMP by human and mouse cells, including cell lines and primary cell *ex vivo*, depends on SLC19A1 expression and function.

**Figure 4.**
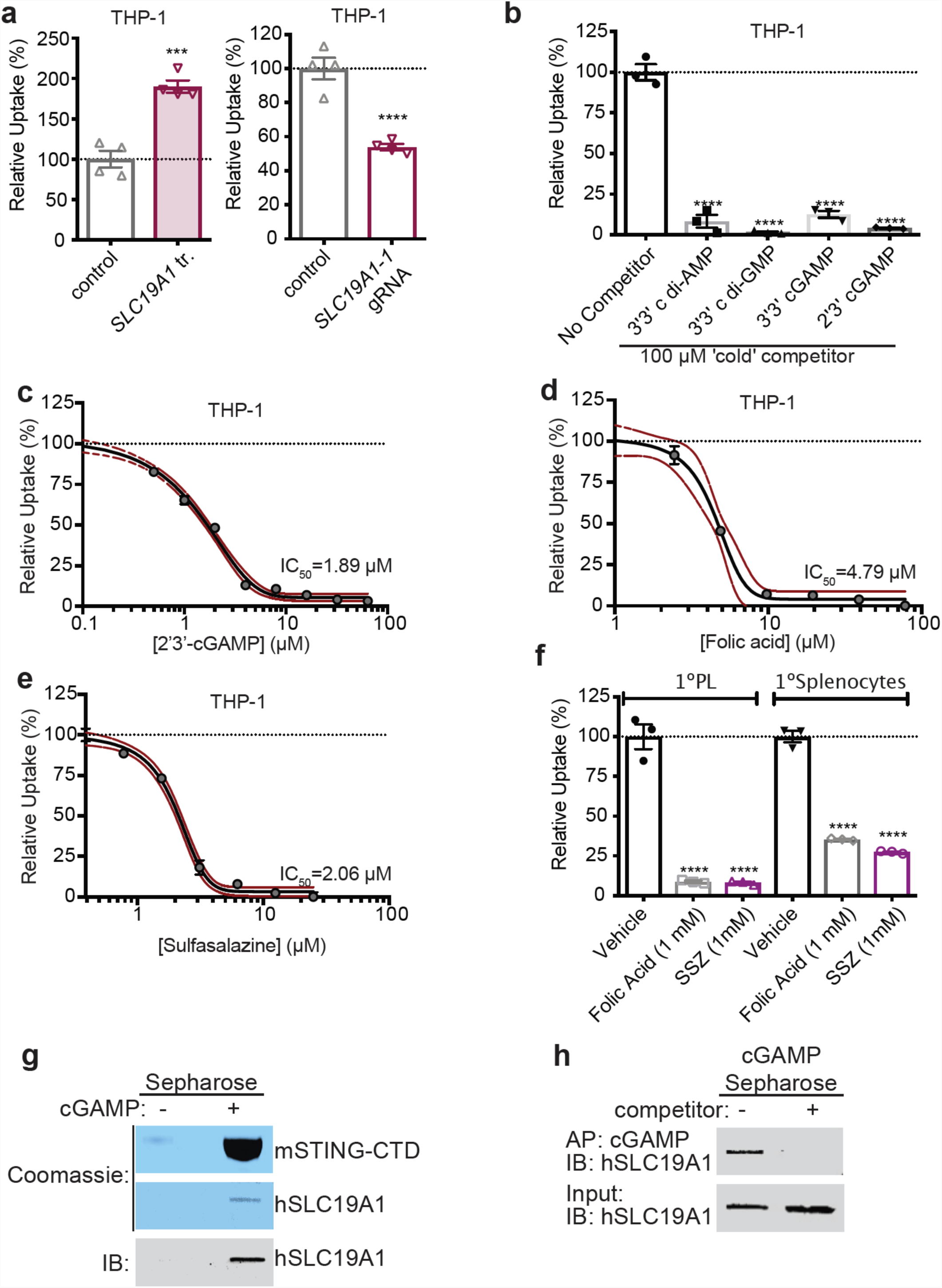
SLC19A1 transports CDNs into cells. **a,** [^32^P] 2’3’-cGAMP uptake by THP-1 monocytes transduced with empty vector (control) or *SLC19A1* expression vector (left panel), or transduced with a non-targeting control CRISPRi gRNA or *SLC19A1* CRISPRi gRNA (right panel). **b,** [^32^P] 2’3’-cGAMP uptake by THP-1 monocytes in the presence of 100 µM competing, unlabeled cyclic di-nucleotides. **c, d**, Competitive inhibition of [^32^P] 2’3’-cGAMP uptake by THP-1 cells in the presence of varying concentrations of competing, unlabeled 2’3’-cGAMP (IC_50_ = 1.89 ± 0.11 µM) or Folic Acid (IC_50_ = 4.79 ± 0.08 µM). **e,** Inhibition of [^32^P] 2’3’-cGAMP uptake by THP-1 cells in the presence of varying concentrations of sulfasalazine (IC_50_ = 2.06 ± 0.17 µM). **f,** [^32^P] 2’3’-cGAMP uptake by primary (1°) peritoneal leukocytes (PL) or splenocytes in the presence of excess folic acid or Sulfasalazine (SSZ). **g,** Binding of SLC19A1 to 2’3’ cGAMP. Coomassie staining and Western blot analysis of pulldowns with 2’3’ cGAMP-Sepharose (+) or control ethanoloamine-Sepharose (−) beads. The beads were incubated with recombinant C-terminal domain of mSTING (mSTING-CTD) or with recombinant hSLC19A1 before precipitation and analysis. **h,** 2’3’cGAMP competes binding of SLC19A1 to 2’3’ cGAMP-Sepharose beads. Soluble 2’3’-cGAMP (250 µM) was added (+) or not (−) to the mixtures of 2’3’ cGAMP-Sepharose and hSLC19A1, before precipitation and Western blot analysis. Data are representative of three independent experiments with similar results. Data are representative of three independent experiments with similar results. In all panels, error bars represent ± SE of biological replicates. Red dashed lines represent the 95% confidence interval for the non-linear regression. Statistical analysis was performed using an unpaired two-tailed Student’s t-test (a) or one-way ANOVA (b and f) followed by Tukey’s post-test. ***P ≤ 0.001; ****P ≤0.0001

If SLC19A1 transports CDNs into cells, CDNs may directly interact with SLC19A1. Consistent with a direct interaction between 2’3’-cGAMP and SLC19A1, His-tagged SLC19A1 was precipitated by 2’3’-cGAMP immobilized on Sepharose beads (Fig. 4g and S8). This interaction was specific, as free, unbound 2’3’-cGAMP competitively disrupted the 2’3’-cGAMP-SLC19A1 interaction (Fig. 4h and S8). As a positive control, His-tagged STING C-terminal domain was also precipitated by 2’3’-cGAMP-Sepharose (Fig 4g and S8). These data suggest that CDNs interact with SLC19A1, consistent with the proposed role of SLC19A1 as a CDN transporter. Taken together, our results demonstrate that SLC19A1 is a mammalian CDN transporter, required for exogenous CDN-mediated type I Interferon activation.

The response to CDNs is weak in most cell lines tested, and can be increased by overexpression of *SLC19A1*. Indeed, THP-1 cells are near the top of a large set of cell lines in expression of both *SLC19A1* and *STING* (Fig. S9), suggesting that *SLC19A1* expression and *STING* expression may together predict the responsiveness to CDN stimulation by cell lines and tumors.

Both folic acid and sulfasalazine almost completely blocked CDN uptake and/or stimulation, whereas CDN stimulation was not completely inhibited in *SLC19A1*-null cells. This implied that another transporter sensitive to folic acid competition and sulfasalazine inhibition may play a role in CDN uptake. Overexpression of *SLC46A1*, which encodes the only other known folate transporter, did increase responses to CDNs (Fig. S10). However, depletion of *SLC46A1* had only a modest effect on CDN stimulation, and was not a significant hit in our screen. Furthermore, depleting *SLC46A1* and *SLC19A1* together was no more effective than depleting SLC19A1 alone (Fig. S11). These data suggest that yet another transporter that is inhibited by folic acid and sulfasalazine may play a partial role in CDN transport. SLC46A3, another transporter, was also a hit in our screen. Overexpression of *SLC46A3* increased the response to CDNs (Fig. S10). Depletion of *SLC46A3* had a modest effect on reporter induction by both CDNs (Fig. S11). However, depleting both *SLC19A1* and *SLC46A3* together did not reduce responses more than depletion of *SLC19A1* alone (Fig. S11), suggesting that SLC46A3 is not responsible for most of the residual CDN transport in *SLC19A1*-depleted cells.

Our findings define SLC19A1 as a major transporter of exogenous 2’3’ cGAMP, 2’3’-RR CDA and probably other CDNs into the cytosol. In this context, it likely plays an important role in the anti-tumor and adjuvant effects of injected CDNs. It may also be important in cell-to-cell transport of CDNs in immune responses, both in the context of cancer^11^ and potentially during viral infections. SLC19A1-mediated uptake of CDNs may also be critical for the pathology of various inflammatory diseases^35,36^. For example, in mouse models of inflammatory bowel disease (IBD), some evidence suggests that host cells import CDNs produced by intestinal bacteria, activating STING in a cGAS-independent fashion^36^. SLC19A1-mediated uptake of CDNs from the extracellular environment may thus contribute to the inflammatory profile underlying such diseases. Moreover, the SLC19A1 inhibitor sulfasalazine is a first line treatment in rheumatoid arthritis, and is often used to treat inflammatory bowel diseases (IBD), including ulcerative colitis and Crohn’s disease^37,38^. Sulfasalazine has an immunosuppressive effect, in part by inhibiting the NF-κB pathway^39^, but the mechanism of inhibition is unknown. Our results raise the intriguing possibility that sulfasalazine exerts its anti-inflammatory effects in these diseases by inhibiting uptake of CDNs produced endogenously or by commensal bacteria, preventing STING activation. In conclusion, we have identified SLC19A1 as a CDN transporter with potential relevance in the context of cancer immunotherapy, immunosurveillance, and inflammatory disease.

## Methods

### Cell culture

All cell lines were cultured at 37°C in humidified atmosphere containing 5% CO2 with media supplemented with 100 U/mL penicillin, 100 µg/mL streptomycin, 0.2 mg/mL glutamine, 10 µg/mL gentamycin sulfate, 20 mM Hepes and 10% FCS. THP-1, C1R, and K562 cells were cultured in RPMI medium, and 293T, 293T transfected with hSTING (293T+hSTING), MDA-MBA-453 (MDA), and RAW macrophages were cultured in DMEM medium. THP-1, K562, 293T cells, and RAW macrophages were present in the lab at the time this study began. MDA cells were obtained from the Berkeley Cell Culture Facility. C1R cells were a generous gift from Veronika Spies (Fred Hutchinson Cancer Center, Seattle WA). 293T+hSTING cells were generated at Aduro Biotech Inc.

### Antibodies and reagents

The following antibodies were derived from Cell signaling technologies: rabbit-anti-human TBK1 mAb (clone D1B4, used 1:500 for immunoblot [IB]), rabbit-anti-human phospho-TBK1 mAb (clone D52C2, 1:1000 for IB), rabbit-anti-human STING mAb (clone D2P2F, 1:2000 for IB), rabbit-anti-human phospho STING mAb (clone D7C3S, used 1:1000 for IB), rabbit-anti-human phospho-IRF3 mAb (clone 4D4G, 1:1000 for IB). Antibodies derived from LI-COR Biosciences: goat-anti-mouse IgG IRDye 680RD conjugated (cat. #: 926-68070, used 1:5000), donkey-anti-rabbit IgG IRDye 800CW conjugated (cat. #: 926-32213), donkey-anti-rabbit IgG IRDye 680RD (cat. #: 926-68073). Other antibodies: rabbit-anti-human IRF3 mAb (Abcam, cat. #: EP2419Y, used 1:2000 for IB), mouse-anti-human transferrin receptor mAb (Thermo Fischer Scientific, clone H68.4, used 1:1000 for IB), rabbit-ant-human SLC19A1 pAb (Picoband, cat. #: PB9504, used 0.4 µg/ml for IB), APC-conjugated mouse-anti-human CD55 mAb (BioLegend, clone JS11, used 1:50 for flow cytometry), mouse-anti-human CD59 mAb (BioLegend clone p282, used 1:250 for flow cytometry), APC-conjugated goat-anti-mouse IgG (BioLegend, cat. #: 405308, used 1:100 for flow cytometry). Reagents used: Sulfasalazine (Sigma-Aldrich, cat. #: S0883), polybrene (EMD Millipore, cat. #: TR1003G), 3’3’-cyclic-di-AMP (CDA) (Invivogen, cat. #: tlrl-nacda), 2’3’-RR CDA and 2’3’-cyclic-di-GMP-AMP (cGAMP) (generous gift from Aduro Bioscience Inc.), human interferon-β (PeproTech, cat. #: 300-02B), mouse interferon-β1 (BioLegend, cat. #: 581302). Antibiotic selection: puromycin (Sigma-Aldrich, cat. #: P8833), blasticidin (Invivogen, cat. #: ant-bl-1, used at 10 µg/ml), zeocin (Invivogen, cat. #: ant-zn-1, used at 200 µg/ml).

### Plasmids

A gBLOCK gene fragment (Integrated DNA Technologies, Inc.) encoding the tdTomato reporter gene driven by the interferon stimulatory response elements (ISREs) and the minimal mouse interferon-β promoter was cloned into a dual promoter lentiviral plasmid by means of Gibson assembly. This lentiviral plasmid co-expressed the Zeocin resistance gene and GFP via a T2A ribosomal skipping sequence controlled by the human EF1A promoter, and was generated as described previously^40^.

For rescue and overexpression of *SLC19A1, SLC46A1*, or *SLC46A3*, a gBLOCK gene fragment encoding *SLC19A1* (gene ID 6573, transcript 1), *SLC46A1* (gene ID 113235) or *SLC46A3* (gene ID 283537) was cloned by Gibson assembly into a dual promoter lentiviral plasmid co-expressing the Blasticidin resistance gene and the fluorescent gene mAmetrine.

For CRISPR interference (CRISPRi)-mediated depletions, cells were transduced with a lentiviral dCas9-HA-BFP-KRAB-NLS expression vector (Addgene plasmid #102244).

For screen validation using individual gRNAs, gRNAs (table S3) were cloned into the same expression plasmid used for the gRNA library (“pCRISPRia-v2”, Addgene plasmid #84832, a gift from Jonathan Weissman). The lentiviral gRNA plasmid co-expresses a puromycin resistance gene and blue fluorescence protein (BFP) via a T2A ribosomal skipping sequence controlled by the human EF1A promoter. The CRISPRi gRNAs introduced into this vector by Gibson assembly were expressed from a murine U6 promoter. For expression of multiple gRNAs, additional gRNAs were introduced in a separate vector that co-expressed the blasticidin resistance gene and mAmetrine via a T2A ribosomal skipping sequence under the control of a human EF1A promoter.

Conventional CRISPR gRNAs (see table S3) were cloned into a selectable lentiviral CRISPR/Cas9 vector. This lentiviral vector includes a human codon-optimized *S.pyogenes* Cas9 co-expressing puromycin resistance gene via a T2A ribosome skipping sequence under the control of a minimal human EF1A promoter^40,41^.

### Lentiviral production and transduction

Lentivirus was produced by transfecting lentiviral plasmids and 2^nd^ generation packaging/polymerase plasmids into 293T cells using TransIT-LT1 Reagent (Mirus Bio LLC). Virus-containing supernatants were harvested 72h later, centrifuged to remove cell debris, and filtered using a 0.45 µm PES filter. Filtered virus supernatant was used to transduce target cells by spin-infection (800 x g for 90min at 33°C) in the presence of 8 µg/ml polybrene. After spin-infection virus and polybrene containing medium was diluted 1:1 with fresh medium. 72 hours after transduction, cells were sorted based on fluorescence expression using a BD FACSAria cell sorter, or selected with relevant selection reagent.

### 2’3’-RR CDA and 2’3’-cGAMP screens

THP-1 cells co-expressing the tdTomato reporter, GFP, and dCas9-BFP were single cell sorted to select for a THP-1 cell clone with efficient dCas9-BFP-knockdown capacity. Clonal populations were transduced with lentiviral vectors encoding gRNAs targeting GFP, *CD55* or *CD59*. After 1 week on puromycin (2 µg/ml) selection, CD55, CD59 and GFP expression were quantified using the BD LSR Fortessa flow cytometer. A clonal cell that showed the highest reduction in all three marker genes was selected for the screens. Two cultures of THP-1 cells were separately transduced with the human genome-wide CRISPRi v2 library^42^. Each library of THP-1 cells was separately screened by treating the cells with 2’3’-RR CDA or 2’3’-cGAMP followed by selection and analysis. Hence, each screen was performed twice, with different CRISPRi library transduced cultures of THP-1 cells.

For each transduction, the THP-1 clone was expanded to 320 million cells and transduced with the human genome-wide CRISPRi v2 library^42^, which contains approximately 100,000 gRNAs targeting around 20,000 genes. Sufficient cells were transduced and propagated to maintain at least 50 million transduced (BFP+) cells, representing 500x coverage of the gRNA library. The transduction efficiency was around 20% to minimize the chance of multiple lentiviral integrations per cell. Two days after transduction, cells were cultured in the presence of puromycin for two days and one day additional day without puromycin. 400 million cells were seeded to a density of 1 million cells/ml and stimulated with 2’3’-RR CDA (2 µg/ml) or 2’3’-cGAMP (15 µg /ml). 20h later, cells were harvested, washed in PBS, and sorted based on BFP expression (presence of gRNAs), GFP expression (presence of reporter) and tdTomato expression using the BD Influx cell sorter and BD FACSaria Fusion cell sorter. The cells were sorted into two populations based on tdTomato expression: the highest 25% of tdTomato expressing cells (hyper-responsive population) and lowest 25% of tdTomato expressing cells (hypo-responsive population). During sorting, all cells were kept at 4°C. After sorting, cells were counted: the sorted populations contained 15-20 million cells, and the unsorted control contained 100-150 million cells. Cells were washed in PBS, and cell pellets were stored at −80°C until further processing.

### gDNA isolation and sequencing

Genomic DNA was isolated from sorted cells using NucleoSpin Blood kits (Macherey-Nagel). PCR was used to amplify gRNA cassettes with Illumina sequencing adapters and indexes as described previously^43^. Genomic DNA samples were first digested for 18 hours with *SbfI*-HF (NEB) to liberate a ∼500 bp fragment containing the gRNA cassette. The gRNA cassette was isolated by gel electrophoresis as described previously^43^. using NucleoSpin Gel and PCR Clean-up kits (Macherey-Nagel), and the DNA was then used for PCR. Custom PCR primers are listed in Supplementary Table 5. Indexed samples were pooled and sequenced on an Illumina HiSeq-2500 for the 2’3’RR CDA screen and an Illumna HiSeq-4000 for the 2’3’-cGAMP screen using a 1:1 mix of two custom sequencing primers (Supplementary Table 5). Sequencing libraries were pooled proportional to the number of sorted cells in each sample. The target sequencing depth was at least 2,000 reads/gRNA in the library for unsorted “background” samples, and at least 10 reads/cell in sorted samples.

### Screen data analysis

CRISPRi samples were analyzed using the Python-based ScreenProcessing pipeline (https://github.com/mhorlbeck/ScreenProcessing). Normalization using a set of negative control genes and calculation of phenotypes and Mann-Whitney *p-value*s was performed as described previously^42,44^. Briefly, Illumina 50bp single end sequencing reads for pooled sublibraries one to four and five to seven were trimmed to 29bp and guides were quantified by counting exact matches to the CRISPRi v2 human library guides. Phenotypes were calculated as the log2 fold change in enrichment of an sgRNA in the high and low samples versus background as well as high versus low, normalized by median subtracting non-targeting sgRNAs^44,45^. Phenotypes from sgRNAs targeting the same gene were collapsed into a single sensitivity phenotype for each gene using the average of the top three scoring sgRNAs (by phenotype absolute value). For genes with multiple independent transcription start sites (TSSs) targeted by the sgRNA libraries, phenotypes and *p-value*s were calculated independently for each TSS and then collapsed to a single score by selecting the TSS with the lowest Mann-Whitney *p-value*. Counts from the ScreenProcessing pipeline were then used as input to the MAGeCK program to obtain FDR scores for filtering (see table S2).

Genes were also ranked by individual gRNAs with the greatest enrichment/depletion between the hypo-responsive and hyper-responsive libraries. gRNA read counts were normalized to library sequencing depth by converting to read counts per million total reads. For each gRNA, the ratio between the read counts for the hypo-responsive and hyper-responsive libraries was found and averaged between replicates. For hypo-responsive gene rankings, each gene was ranked by the single corresponding gRNA with the highest hypo-to-hyper ratio (see table S1, ‘highest ratio hypo/hyper’ column). For hyper-responsive gene rankings, each gene was ranked by the single corresponding gRNA with the lowest hypo-to-hyper ratio (see table S1, ‘lowest ratio hypo/hyper’ column). Gene-level phenotypes are available as Supplemental Materials (table S1 and S2).

### CDN and IFN-β Stimulation assays

The week prior to stimulation experiments, cells were cultured at the same density. The day before stimulation, cells were seeded to 0.5×10^5^ cells/ml. Cells were stimulated with CDNs or IFN-β in 48W plates using 50,000 cells/well in 300 µl medium. After 18-24h, cells were transferred to a 96W plate and tdTomato expression was measured by flow cytometry using a high throughput plate reader on a BD LSR Fortessa. For stimulations in the presence of sulfasalazine, cells were stimulated in 48W plates using 20,000 cells/well in 300 µl medium. Cells were incubated with sulfasalazine or DMSO as vehicle prior to stimulations with CDNs or IFN-β. 18-24h after stimulation, tdTomato reporter expression was quantified by flow cytometry using a high throughput plate reader on a BD LSR Fortessa.

### Production of SLC19A1 knockout cell lines

THP-1 cells expressing the tdTomato reporter were transduced with a CRISPR/Cas9 lentiviral plasmid encoding a control gRNA or a gRNA targeting *SLC19A1* at a region critical for transport^46^ (see table S3). Transduced cells were selected using puromycin for 2 days and single cell sorted using a BD FACSAria cell sorter. Control cells and *SLC19A1*-targeted cells were selected that had comparable forward and side scatter by flow cytometry analysis. Genomic DNA was isolated from clones using the Qiamp DNA minikit (Qiagen), and the genomic region surrounding the *SLC19A1* gRNA target site was amplified by PCR using primers 5’-TTCTCCACGCTCAACTACATCTC-3’ and 5’-CAGCATCCGCGCCAGCACTGAGT-3’. PCR product was cloned into 5-alpha competent bacteria (New England Biolabs, cat. #C2987) using a TOPO TA cloning kit (Thermo Fischer Scientific, cat. # 450641) according to manufacturer’s instructions. After blue/white screening, a minimum of 10 colonies were sequenced per THP-1 clone, and sequences were analyzed using SeqMan (Lasergene DNASTAR). THP-1 clones with out-of-frame mutations at the SLC19A1 gRNA target site were selected for further experiments.

### RT-qPCR

Cells were harvested and washed in ice-cold PBS. Cells were transferred to RNase-free microcentrifuge tubes and RNA was isolated using the RNeasy mini kit (Qiagen, cat. #: 74104) including a DNase step (Qiagen, cat. #: 79254). RNA concentration was measured by NanoDrop (Thermo Fischer), and 1 µg of RNA was used as input for cDNA synthesis using the iScript cDNA synthesis kit (Bio-rad, cat. #: 1708890). cDNA was diluted to 20 ng/µl and 2.5 µl/reaction was used as input for the qPCR reaction. qPCR reactions were set up using SSOFast EvaGreen Supermix (Bio-Rad, cat. #: 1725200) according to the manufacturer’s recommendations, using 500 nM of each primer and following cycling conditions on a Bio-Rad C1000 Thermal Cycler: 2 min at 98°C, 40 repeats of 2 sec at 98°C and 5 sec at 55°C. Primers used to amplify the *HPRT1, YHWAZ, CCL5, CXCL10, STING, IRF3, SLC19A1, SLC46A1*, and *SLC46A3*-specific PCR products are listed in Table S4. The housekeeping genes *HPRT1* and *YHWAZ* served as endogenous control.

For quantification of *IFNB1* mRNA, RNA was extracted with the Nucleospin RNA Isolation Kit (Machery-Nagel) and reverse-transcribed with the iScript cDNA synthesis kit (Bio-Rad). TaqMan real-time qPCR assays were used for quantification of human *IFNB1* (Hs01077958_s1). *ACTB* (Hs01060665_g1) served as an endogenous control.

### Synthesis of [^32^P] cyclic GMP-AMP and [^32^P] cyclic di-AMP

Radiolabeled 2’3’ cGAMP was enzymatically synthesized by incubating 0.33 µM α-[^32^P] ATP (Perkin-Elmer) with 250 µM unlabeled GTP, 1 µg of Interferon Stimulatory DNA 100mer (kindly provided by Daniel Stetson), and 1 µM of recombinant His-tagged 2’3’ cGAMP Synthase (cGAS) in binding buffer [40 mM Tris pH 7.5, 100 mM NaCl, 20 mM MgCl_2_] at 37°C overnight. The reaction was confirmed to have gone to completion by Thin Layer Chromatography (TLC) analysis. Briefly, the 2’3’ cGAMP synthesis reaction was separated on Polygram CEL300 PEI TLC plates (Machery-Nagel) in buffer containing 1:1.5 (vol/vol) saturated (NH_4_)_2_SO_4_ and 1.5 M NaH_2_PO_4_ pH 3.6. The TLC plates were then air dried and exposed to a PhosphorImager screen for visualization using a Typhoon scanner (GE Healthcare Life Sciences). Next, the sample was incubated with HisPur Ni-NTA resin (Thermo Scientific) for 30 min in order to remove recombinant cGAS. The resultant slurry was transferred to a minispin column (Thermo Scientific) to elute crude [^32^P] 2’3’ cGAMP. Recombinant mSTING-CTD protein was used for further purification of synthesized [^32^P] 2’3’ cGAMP. 100 µM mSTING-CTD was bound to HisPur Ni-NTA resin and incubated with the remaining crude 2’3’ cGAMP synthesis reaction mixture for 30 min on ice. Following removal of the supernatant, the Ni-NTA resin was washed three times with cold binding buffer. The resin was then incubated with 100 µL of binding buffer for 10 min at 95 °C, and transferred to a minispin column to elute [^32^P] 2’3’ cGAMP. The resulting STING-purified [^32^P] 2’3’ cGAMP was evaluated by TLC analysis and determined to be ∼99% pure.

Radiolabeled c di-AMP was synthesized as described previously ^47^. Briefly, 1 µM α-[^32^P] ATP (Perkin-Elmer) was incubated with 1 µM of recombinant DisA in binding buffer at 37°C overnight. The reaction mixture was boiled for 5 min at 95°C and DisA was removed by centrifugation. Recombinant His-tagged RECON was then used to further purify the c di-AMP reaction mixture. 100 µM His-tagged RECON was bound to HisPur Ni-NTA resin for 30 min on ice. The resin was washed three times with cold binding buffer and then incubated with 100 µL of binding buffer for 5 min at 95°C. The slurry was then transferred to a minispin column to elute [^32^P] c di-AMP. The purity of the radiolabeled c di-AMP was assessed by TLC and determined to be ∼98%.

### Nucleotide-Binding Assays

The ability of radiolabeled 2’3’ cGAMP and c di-AMP to bind recombinant STING was evaluated by DRaCALA (differential radial capillary action of ligand assay) analysis, as previously described^48^. Briefly, varying concentrations of recombinant STING were incubated with ∼1 nM of radiolabeled cyclic di-nucleotide in binding buffer for 10 min at room temperature. The reaction mixtures were blotted on nitrocellulose membranes and air dried for 15 min. The membranes were then exposed to a PhosphorImager screen and visualized using a Typhoon scanner.

### Nucleotide-Uptake Assays

For transport assays, cells were collected by centrifugation and washed in Dulbecco’s Phosphate-Buffered Saline (DPBS) (Life Technologies). The cell pellets were re-suspended in pre-warmed RPMI 1640 medium (GIBCO) containing 10% heat-inactivated FBS (HyClone) and supplemented with 10 mM HEPES, 1 mM sodium pyruvate and 2 mM L-Glutamine (Thermo Fisher) to a final cell density of 1 × 10^7^ cells per ml. Uptake of 1 nM [^32^P] cGAMP and c di-AMP was assayed in cell suspensions at 37°C over the indicated time points. At the end of each time point, transport was quenched by the addition of cold DPBS. Cells were washed three times with cold DPBS, followed by lysis in 50 µL of cold deionized water. The cell lysates were then transferred to 5 ml of liquid scintillation cocktail (National Diagnostics) and the associated radioactivity was measured by liquid scintillation counting using a LS6500 Liquid Scintillation Counter (Beckman Coulter). For each sample, [^32^P] cyclic di-nucleotide uptake (counts per minute) was normalized to cell count. For competition experiments, cells were pre-incubated with indicated concentrations of “cold” unlabeled ligand for 15 minutes prior to the addition of 1 nM “hot” [^32^P] cGAMP. Cells were then collected at the indicated time points and processed as described above.

### Protein Expression and Purification

Full-length human SLC19A1 cDNA with a C-terminal 8 X His-tag was subcloned into a dual promoter lentiviral vector (see above). Recombinant His-tagged SLC19A1 was expressed using a FreeStyle 293 Expression System. Briefly, 293F cells (1 × 10^6^ cells per ml) grown in FreeStyle 293 Media supplemented with GlutaMax (GIBCO) were transfected with the SLC19A1 expression construct (1µg plasmid DNA per ml of cells) using PEI transfection reagent. Transfected cells were grown for 72 hours in a shaking incubator at 37°C in 5% CO_2_. Three days after transfection, the cells were harvested by centrifugation and washed in DBPS. Cell pellets were then re-suspended in lysis buffer [25 mM Tris pH 8.0, 150 mM NaCl, 1 mM phenylmethylsulfonyl fluoride] supplemented with HALT Protease and Phosphatase Inhibitor Cocktail (Thermo Scientific) and lysed by sonication. The cell lysate was supplemented with 2% (w/v) n-dodecyl-β-D-maltoside (DDM) and rotated for 2h at 4°C. The cell lysates were centrifuged at 15,000 rpm for 1h at 4°C to remove cell debris, and the detergent-soluble fraction was incubated with HisPur Ni-NTA resin for 1h at 4°C. The resin was washed with 100 column volumes of wash buffer [25 mM Tris pH 6.0, 150 mM NaCl, 30 mM imidazole, 5% glycerol (v/v), and 0.05% DDM (w/v)], and bound proteins were eluted in elution buffer [25 mM Tris pH 6.0, 150 mM NaCl, 300 mM imidazole, 5% glycerol (v/v), and 0.05% DDM (w/v)]. The resulting proteins were analyzed by SDS-PAGE followed by Coomasie staining and immunoblotting to confirm expression and purification of His-tagged SLC19A1.

Recombinant cGAS, DisA, mSTING-CTD, and mRECON were expressed and purified as previously described^47–49^. Briefly, plasmids for cGAS, DisA, mSTING-CTD, and mRECON expression were transformed into Rosetta (DE3) pLysS chemically competent cells. Overnight cultures of the resulting transformed bacteria were inoculated into 1.5 L of LB broth at a 1:100 dilution. Bacterial cultures were grown at 37°C to OD_600_ 0.5 followed by overnight induction at 18°C with 0.5 mM isopropyl β-D-1-thiogalactopyranoside (IPTG). Cells were harvested and lysed in PBS supplemented with 1 mM PMSF and soluble protein was purified using nickel-affinity chromatography followed by gel filtration chromatography (S-300, GE Healthcare, Piscataway, New Jersey, USA). After SDS-PAGE analysis, the purified proteins were concentrated in storage buffer [40 mM Tris pH 7.5, 100 mM NaCl, 20 mM MgCl_2_, 25% glycerol (v/v)] and stored at −80°C.

### Synthesis of cGAMP Sepharose

2’3’ cyclic GMP-AMP was enzymatically synthesized using recombinant cGAS as described previously^8,48^. Approximately, 100 mg of purified cGAMP was dissolved in PBS to 200 µM. The pH of the solution was adjusted to 7.5 with NaOH, and the resulting solution was added directly to washed epoxy-activated Sepharose and incubated at 56°C for 2 days. The Sepharose was washed and the absorbance spectrum of 50% slurry was measured to ensure nucleotide coupling. HPLC analysis of the remaining uncoupled nucleotide ensured no degradation of cGAMP occurred during the 2-day incubation. The remaining epoxy groups were blocked with ethanolamine following the instructions provided by GE. In parallel with this blocking step, fresh epoxy-activated Sepharose was also treated with ethanolamine to generate control resin.

### cGAMP Pulldowns

Following nickel affinity purification, recombinant His-tagged SLC19A1 was incubated with 100 µL of ethanolamine-or cGAMP-conjugated Sepharose beads for 4h at 4°C with rotation, as described previously (Sureka et. al., 2014; McFarland et. al., 2016). Beads were washed three times with wash buffer [25 mM Tris pH 6.0, 150 mM NaCl, 5% glycerol (v/v), and 0.05% DDM (w/v)], and bound proteins were eluted by boiling in SDS-PAGE sample loading buffer for 5 min at 95°C. The soluble fraction was then removed and analyzed by SDS-PAGE followed by Coomassie Blue staining and immunoblotting. As a control, recombinant His-tagged mSTING-CTD was incubated with ethanolamine-or cGAMP-conjugated sepharose beads, as described above. Beads were washed three times with binding buffer, and then boiled in SDS-PAGE sample loading buffer for 5 min at 95°C. The soluble fraction was then analyzed by SDS-PAGE followed by Coomasie staining.

### Cell lysis and immunoblotting

For anti-SLC19A1 immunoblotting, cells were lysed and proteins were separated by SDS-PAGE as described above in the ‘cGAMP pulldowns’ paragraph. SDS-PAGE-separated proteins were transferred onto nitrocellulose membranes (Bio-Rad) at 30V overnight at 4°C. Membranes were then air dried for 1h and blocked in 5% Blotto, non-fat milk (NFM, Santa Cruz Biotechnology) in 1 X TBS. Membranes were probed in 5% Bovine Serum Albumin (Fisher) in 1 X TBS-T with anti-SLC19A1 Picoband antibody (Boster Bio).

For protein detection using all other antibodies, cells were counted, washed with PBS and lysed in RIPA buffer (25 mM Tris-HCl pH 7.5, 150 mM NaCl, 1 mM EDTA, 1% NP-40, 0.1% SDS) including cOmplete ULTRA protease inhibitors (Sigma-Aldrich cat. #: 05892791001), phosphatase inhibitors (Biomake, cat. # B15001) and 50mM DTT. Cells lysates were mixed with 4x NuPage LDS sample buffer (Invitrogen cat. #: NP0007), pulse sonicated and incubated at 75°C for 5min. Lysates were loaded onto Bolt 4-12% Bis-Tris Plus SDS-PAGE gels (Invitrogen cat. #: NW04125BOX). SDS-PAGE separated proteins were transferred onto Immobilon-FL PVDF membranes (EMD Millipore) at 100V for 1h at 4°C. Membranes were blocked in 4% NFM, and probed in 1% NFM overnight at 4°C with primary antibody. Membranes were subsequently washed 3 times in 1x-TBS-T and probed with secondary antibody for 1h at RT protected from light. Membranes were washed 2 times in TBS-T, once in TBS, and blots were imaged using an Odyssey CLx System (LI-COR).

### Mice

C57BL/6J mice were obtained from The Jackson Laboratory. All of the mice were maintained in specific pathogen free conditions by the Department of Comparative Medicine at the University of Washington School of Medicine. All experimental procedures using mice were approved by the Institutional Animal Care and Use Committee of the University of Washington and were conducted in accordance with institutionally approved protocols and guidelines for animal care and use.

### Isolation of Mouse Peritoneal Cavity Cells and Splenocytes

Mouse peritoneal cavity cells were recovered by peritoneal lavage with 5 ml ice cold PBS supplemented with 3% FCS, as previously described^50^. The peritoneal cells were cultured in RPMI 1640 medium (GIBCO) supplemented with 10% (v/v) heat-inactivated FBS (HyClone), 10 mM HEPES, 1 mM sodium pyruvate, 2 mM L-Glutamine (Thermo Fisher), 100 U/ml penicillin, 100 µg/ml streptomycin at 37°C in the presence of 5% CO_2_.

For isolation of murine splenocytes, spleens were removed from mice, strained through a 70 µm cell strainer, and homogenized into a single cell suspension using ice cold PBS supplemented with 3% FCS. Red blood cells were lysed by resuspending spleen cells in Red Blood Cell Lysing Buffer (Sigma) and incubating on ice for 10 min. Splenocytes were washed, resuspended in RPMI 1640 medium (GIBCO) supplemented with 10% (v/v) heat-inactivated FBS (HyClone), 10 mM HEPES, 1 mM sodium pyruvate, 2 mM L-Glutamine (Thermo Fisher), 100 U/ml penicillin, 100 µg/ml streptomycin, and used immediately for [^32^P] cGAMP uptake assays.

## Data availability

Raw sequencing data from the CRISPRi screen will be deposited to NCBI GEO prior to final publication.

### Acknowledgements

We thank Lily Zhang and Erik Seidel for lab and technical assistance, Hector Nolla and Alma Valeros for assistance with cell sorting, the UC Berkeley High Throughput Screening Facility for preparation of gRNA lentivirus, Adelle P. McFarland for assistance in the isolation of primary cells from mice, Shana L McDevitt for assistance with deep-sequencing, and Raulet lab members, Russell Vance, Michel DuPage, Jeremy Thorner and Andrea Van Elsas for helpful discussions. RDL is supported by a Cancer Research Institute Irvington Postdoctoral Fellowship. DHR is supported by NIH grants R01-AI113041 and R01-CA093678. BG is supported by the IGI-AstraZeneca Postdoctoral Fellowship, JJW is supported by the Pew Scholars Program in the Biomedical Sciences and 1R21AI137758-01. SAZ is supported by grants from the University of Washington/Fred Hutchinson Cancer Research Center Viral Pathogenesis Training Program (AI083203), the University of Washington Medical Scientist Training Program (GM007266), as well as the Seattle ARCS foundation. JEC is supported by the National Institute Health New Innovator Awards (DP2 HL141006), the Li Ka Shing Foundation and the Heritage Medical Research Institute.

This work used the Vincent J. Coates Genomics Sequencing Laboratory at UC Berkeley, supported by NIH S10 Instrumentation Grants S10 OD018174, S10RR029668 and S10RR027303.

## Author contributions

RDL, SAZ, and NG performed and analyzed the experiments, LO, SMM, and GEK assisted with the experiments, SW and BGG analyzed the deep-sequencing data and advised on the screen design, RDL, SAZ, BGG, JEC, JW, and DHR designed the experiments, RDL, SAZ, JJW, and DHR prepared the manuscript. All authors critically read the manuscript.

## Supplemental information

Table S1. Ranking of target genes based on the ratio between individual gRNAs present in the populations that were hyper-responsive (hyper) or hypo-responsive to CDN treatment (included as Excel file).

Table S2. Ranking of targeted genes present in the populations hyper-responsive (hyper) or hypo-responsive (hypo) to CDN treatment. RRA ranking is based on the score computed by the MaGeCK program, and phenotypes and p-value calculated by the ScreenProcessing pipeline. (included as Excel file)

**Table S3.**
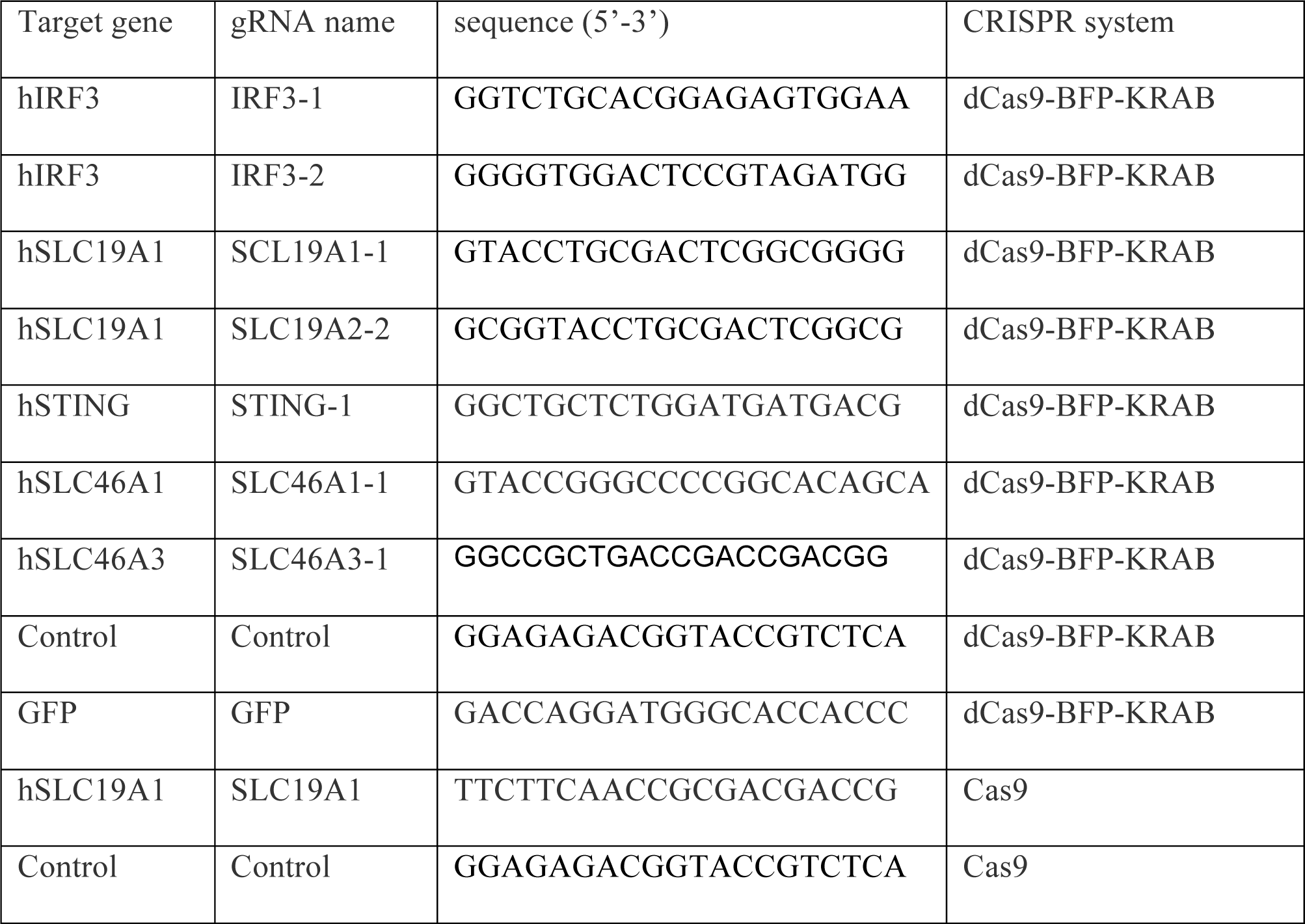
guide RNAs

**Table S4.**
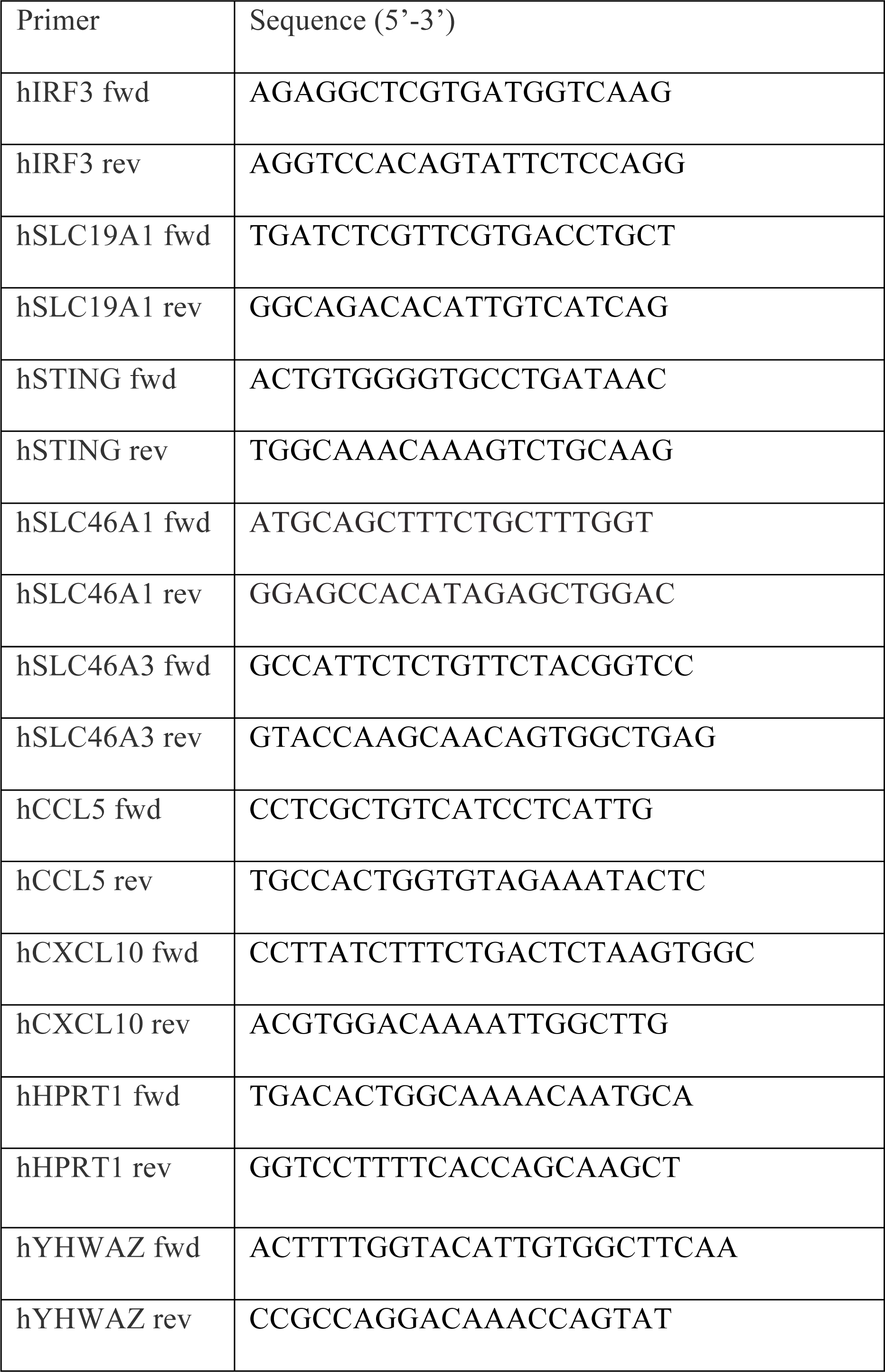
qPCR primers

Table S5. Primers and conditions used for the CDN screens.

**Figure S1.**
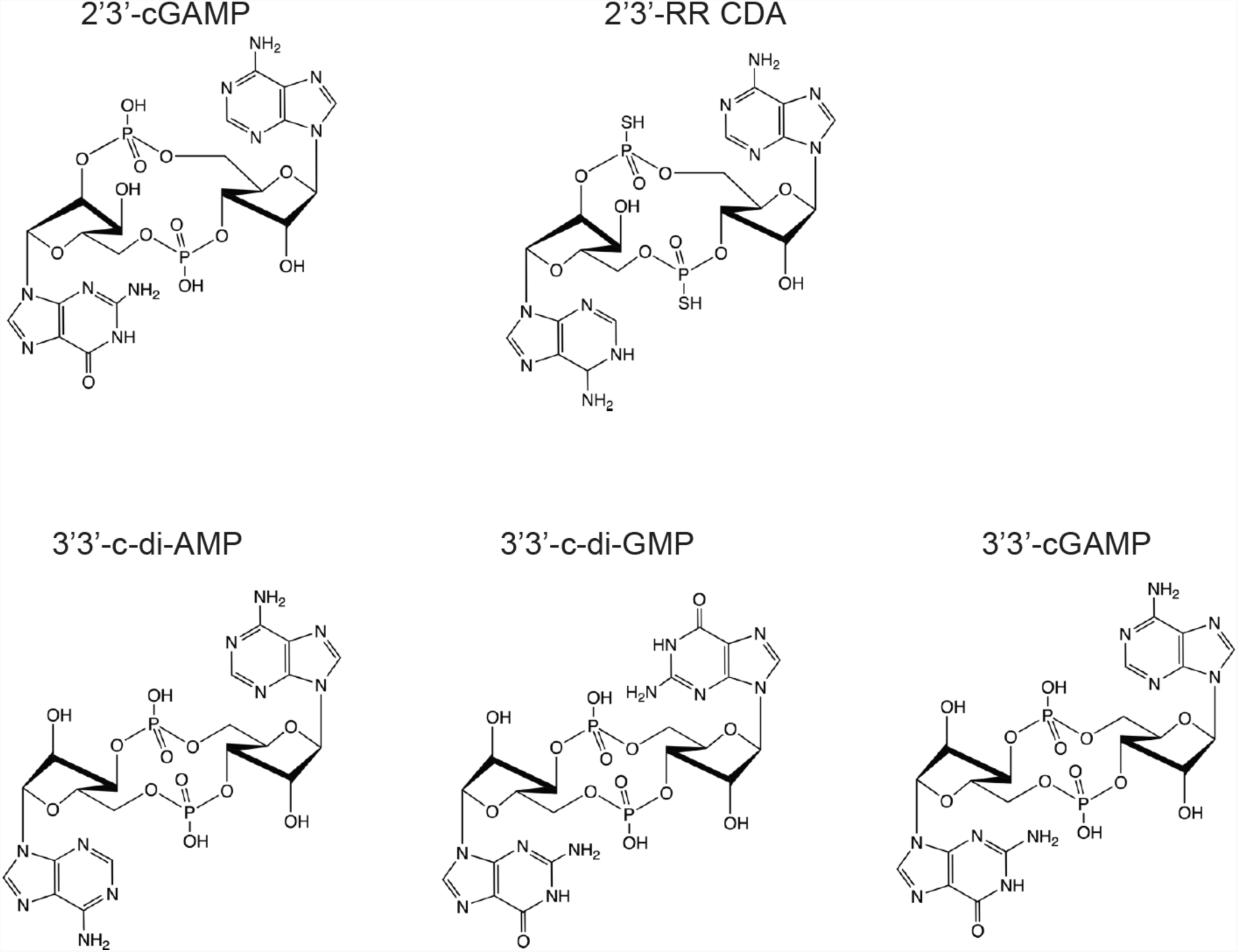
Structures of the CDNs used in this study.

**Figure S2.**
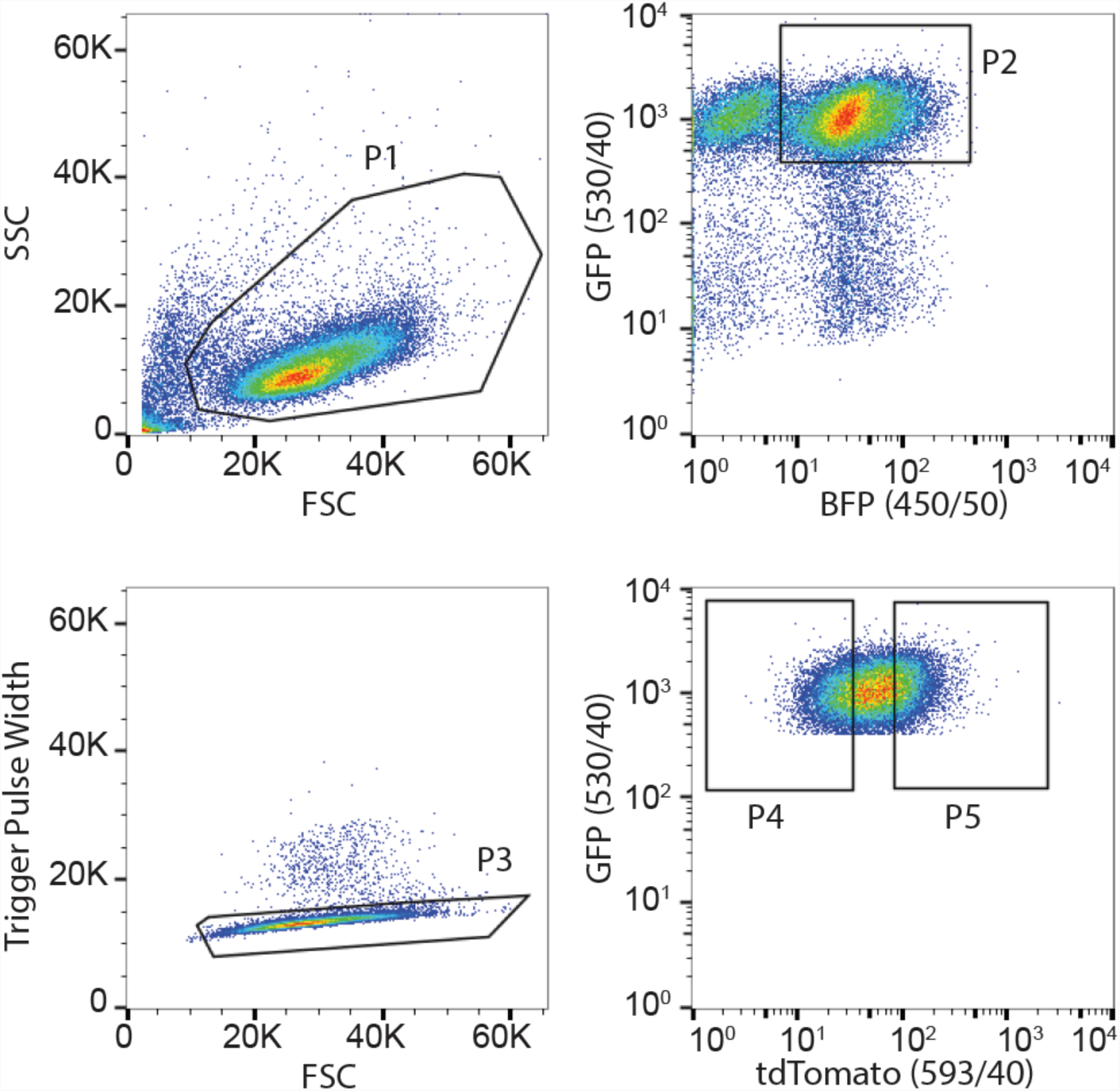
Representative gating strategy for flow cytometry based sorting of the CRISPRi library of reporter-expressing THP-1 cells stimulated with CDNs. Cells were gated based on their forward scatter (FSC) and side scatter (SSC) using gate P1. P1-population was selected based on the expression of blue fluorescent protein (BFP, fluorescent marker for the CRISPRi gRNAs) and GFP (marker for the expression of the reporter construct) using gate P2. In gate P3, the doublet cells present in gate P2 were excluded. In gate P4 and P5, population P3 was gated based on tdTomato expression. The lowest 25% of cells expressing tdTomato were selected in gate P4, and the highest 25% of cells expressing tdTomato were selected in gate P5.

**Figure S3.**
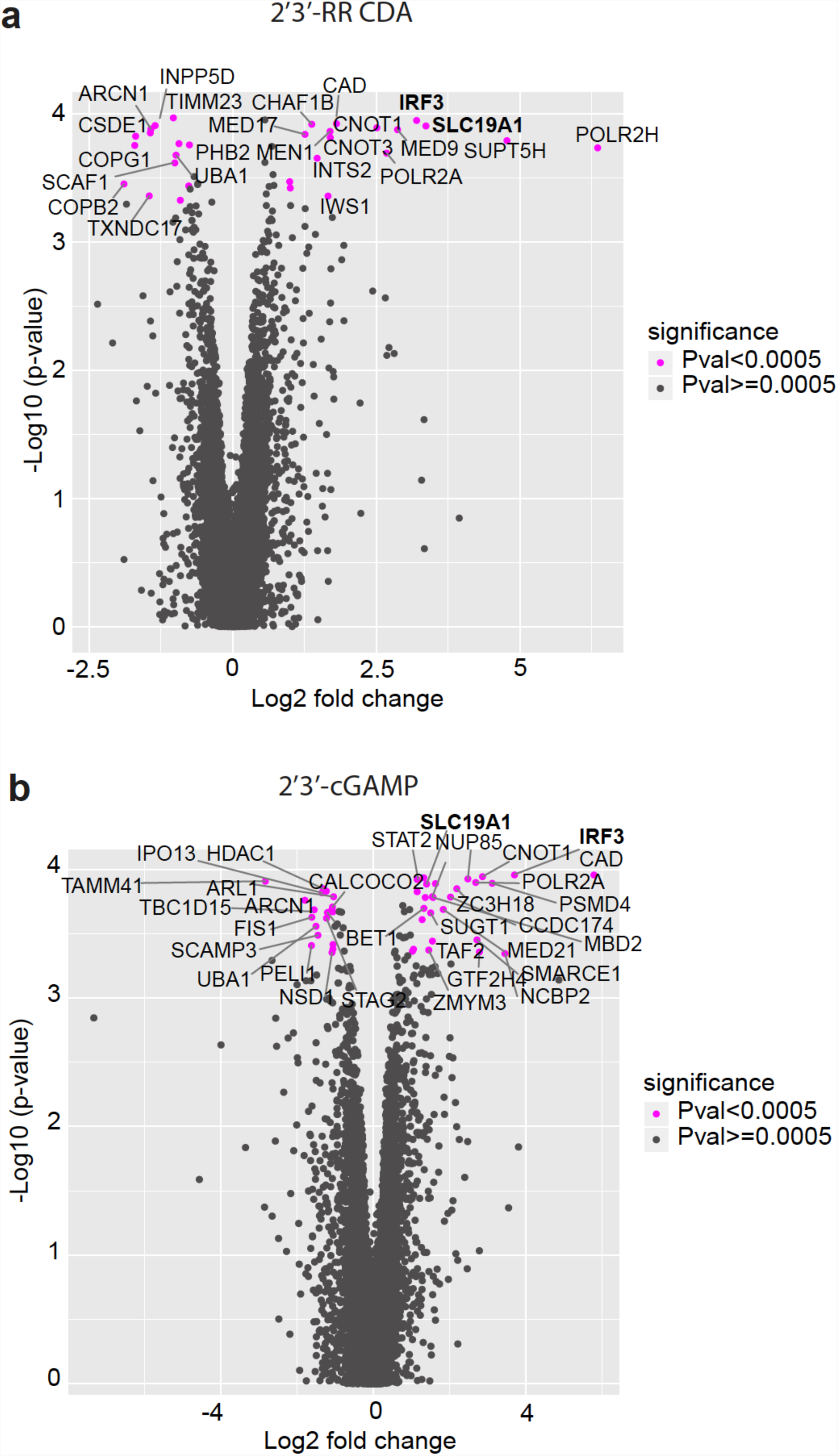
Results of genome-wide CRISPRi screen for host factors crucial for cyclic dinucleotide (CDN) stimulation. Volcano plots of the gRNA-targeted genes enriched or depleted in the tdTomato reporter-low versus reporter-high groups after stimulation with (a) 2’3’-RR CDA or (b) 2’3’-cGAMP. FC: fold change. Each panel represent the combined results of two independent screens.

**Figure S4.**
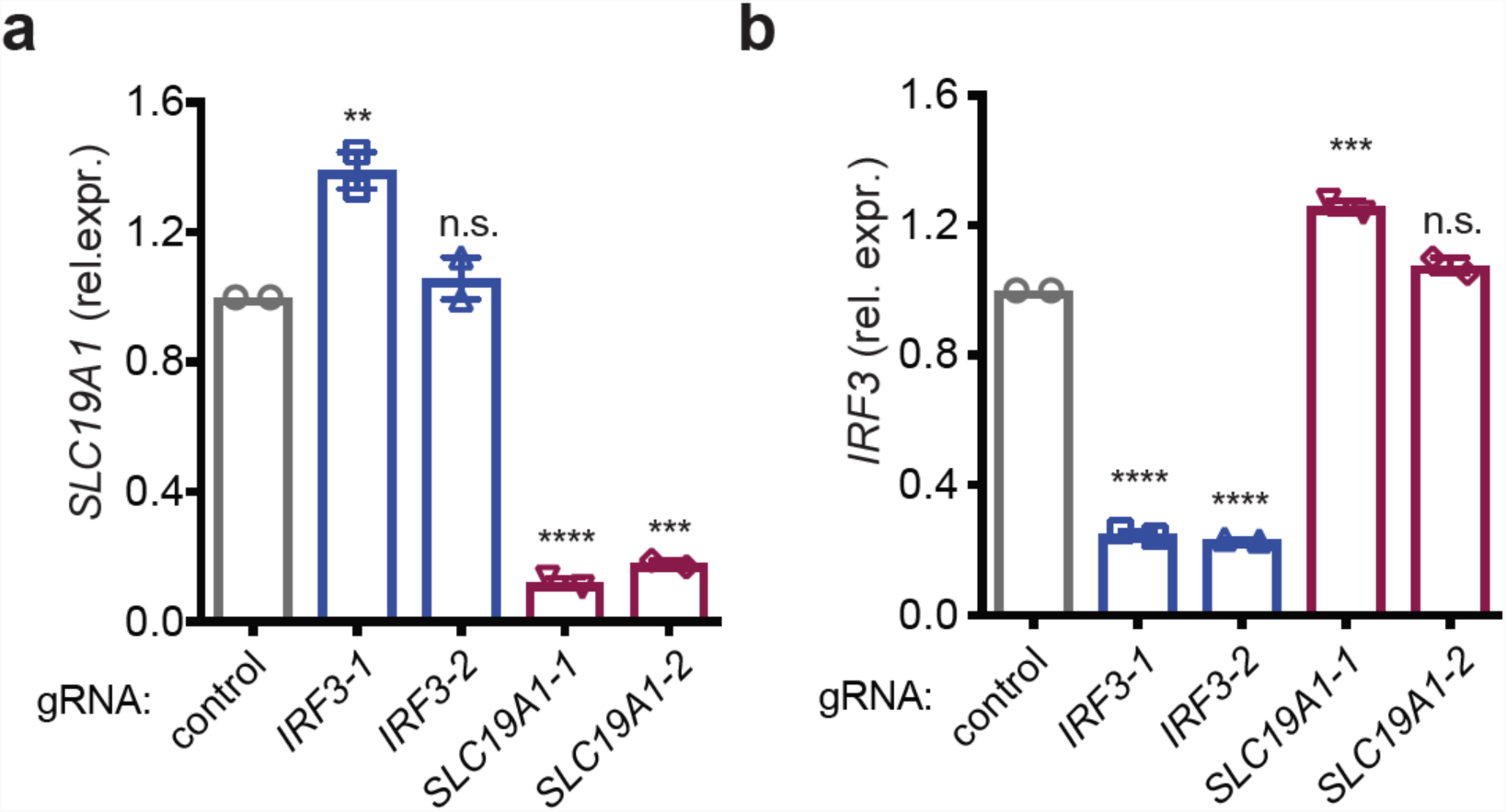
SLC19A1 is critical for CDN-induced reporter expression. a,b, mRNA expression levels of (a) *SLC19A1* or (b) *IRF3* in THP-1 cells expressing a CRISPRi vector and a control non-targeting gRNA or gRNAs targeting IRF3 or SLC19A1 (two gRNAs each). Error bars represent ± SE of at two biological replicates. Statistical analysis was performed to compare each cell line to the control using a one-way ANOVA followed by Dunnetts’s post-test. \*\*\**P* ≤0.001; \*\*\*\**P* ≤ 0.0001; n.s. not significant.

**Figure S5.**
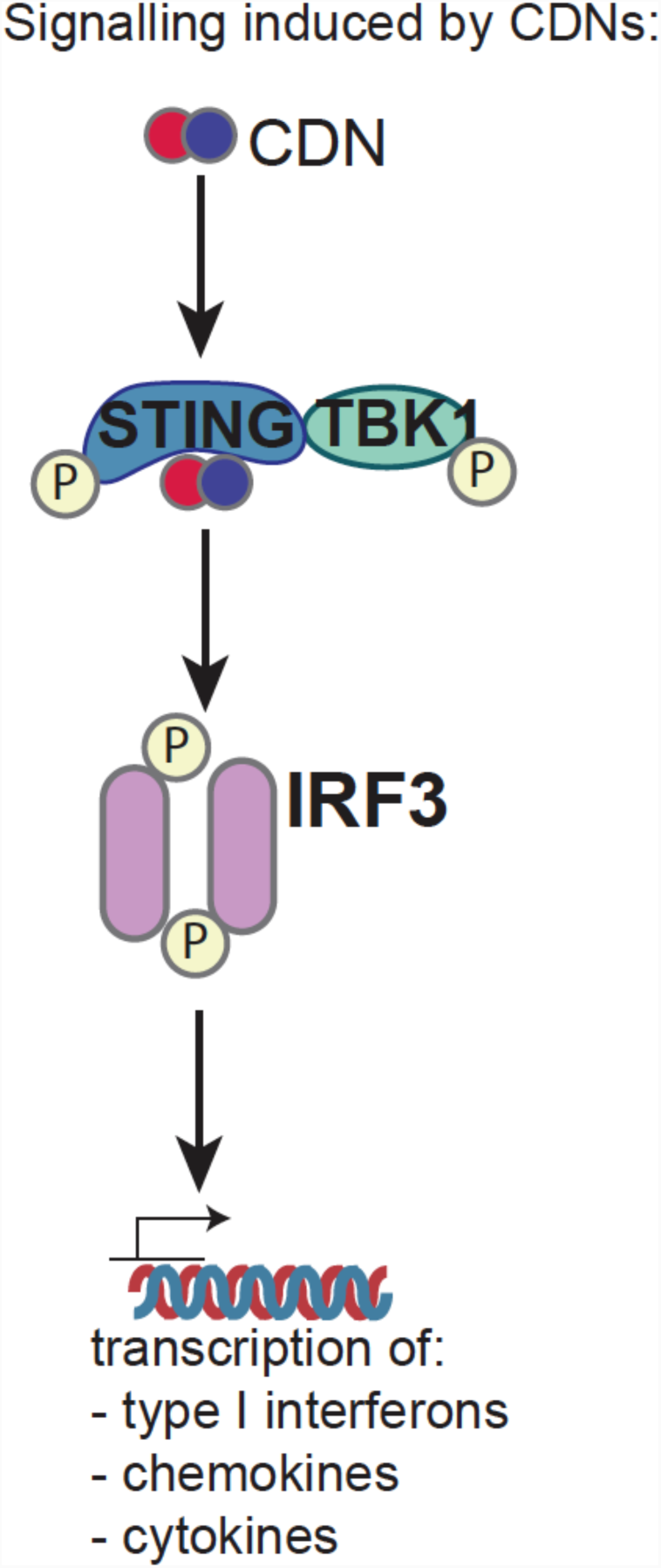
Schematic overview of CDN-induced phosphorylation (P) of STING and downstream effectors TBK1 and IRF3.

**Figure S6.**
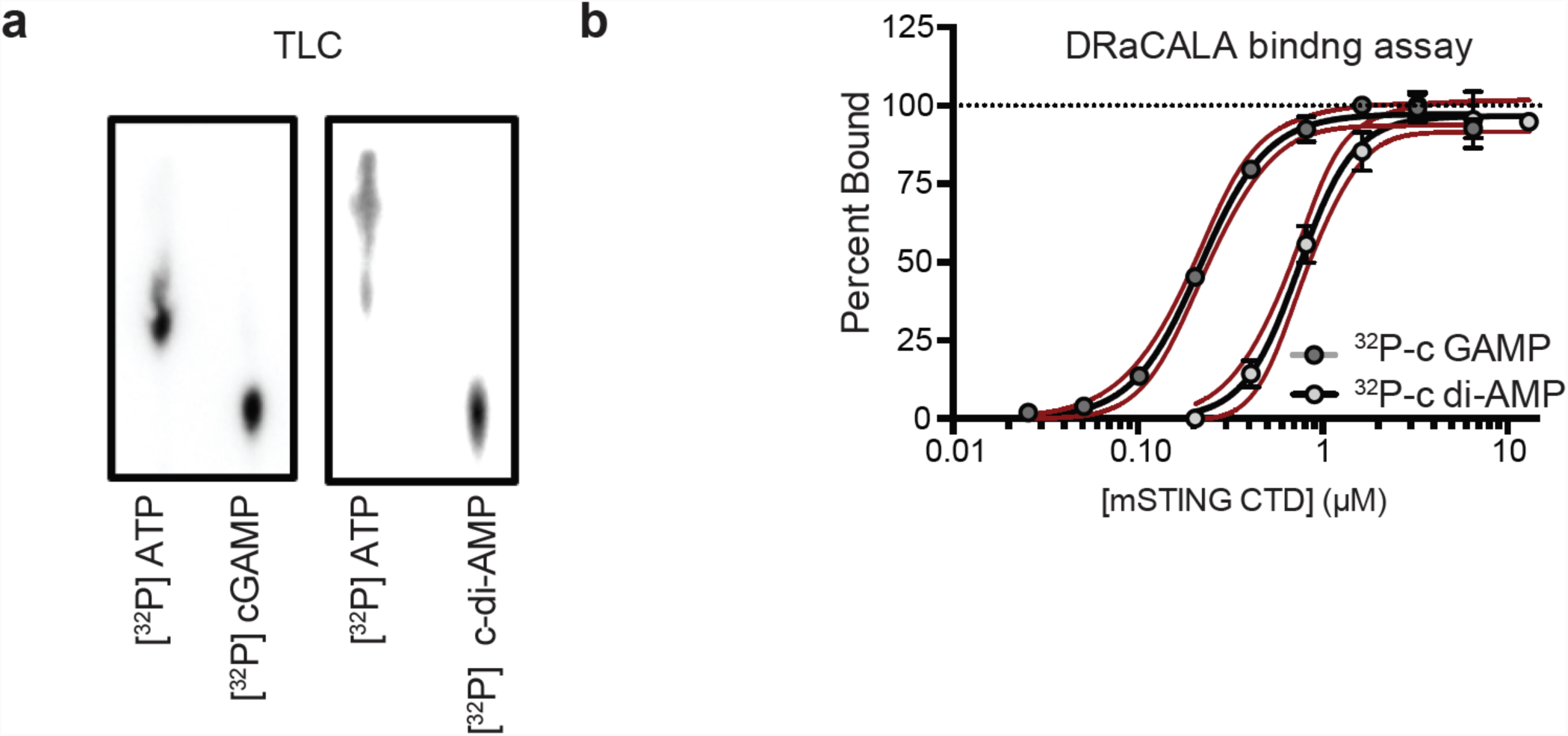
Analysis of enzymatically generated [^32^P] cyclic dinucleotides. **a,** Thin layer chromatography (TLC) analysis of [^32^P] ATP and enzymatically synthesized [^32^P] 2’3’ cGAMP and c di-AMP. **b,** Binding titration of [^32^P] 2’3’ cGAMP or c-di-AMP with mSTING C-Terminal Domain (CTD), determined with DRaCALA assays. Red dashed lines represent the 95% confidence interval for the non-linear regression.

**Figure S7.**
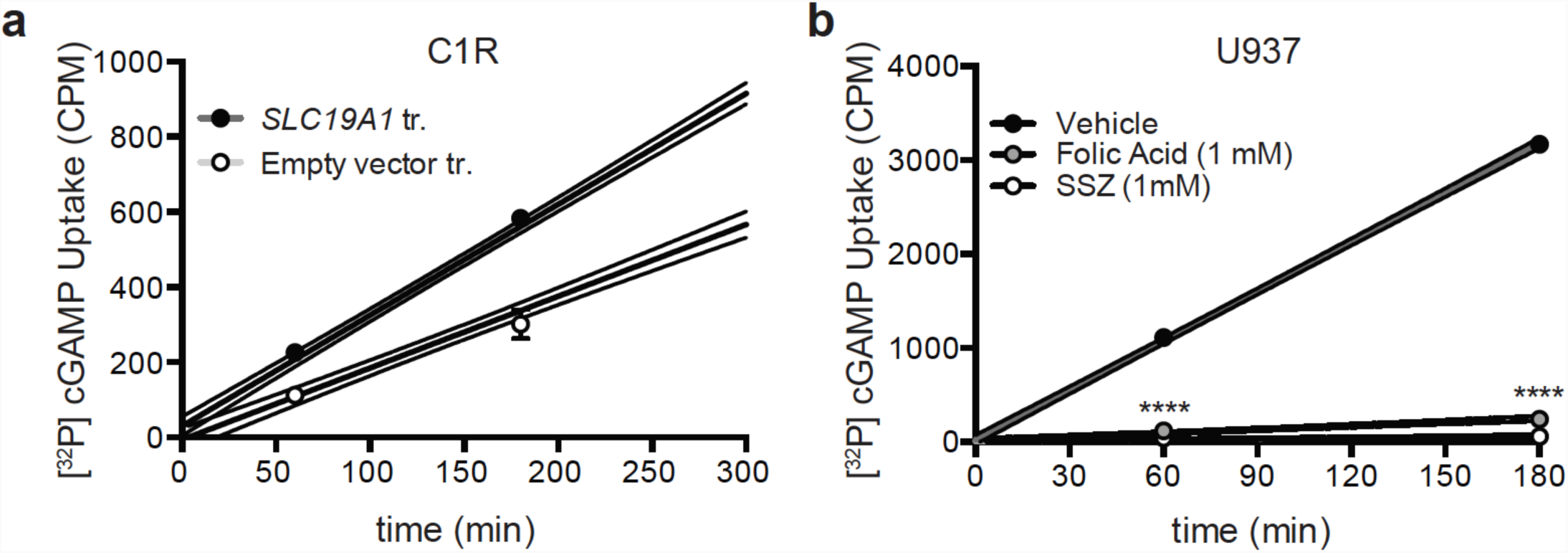
CDN uptake in C1R and U937 cells. **a,** Time course of [^32^P] 2’3’ cGAMP uptake by CIR cells transduced (tr.) with empty vector or *SLC19A1*. **b,** Time course of [^32^P] 2’3’ cGAMP uptake by U937 monocytes in the presence of excess folic acid or sulfasalazine (SSZ). In all panels, error bars represent ± SD of biological replicates. Dashed lines represent the 95% confidence interval for the non-linear regression. Statistical analysis was performed using an unpaired two-tailed Student’s t-test (a) or one-way ANOVA (b) followed by Tukey’s post-test. ***P ≤ 0.001; ****P ≤ 0.0001

**Figure S8.**
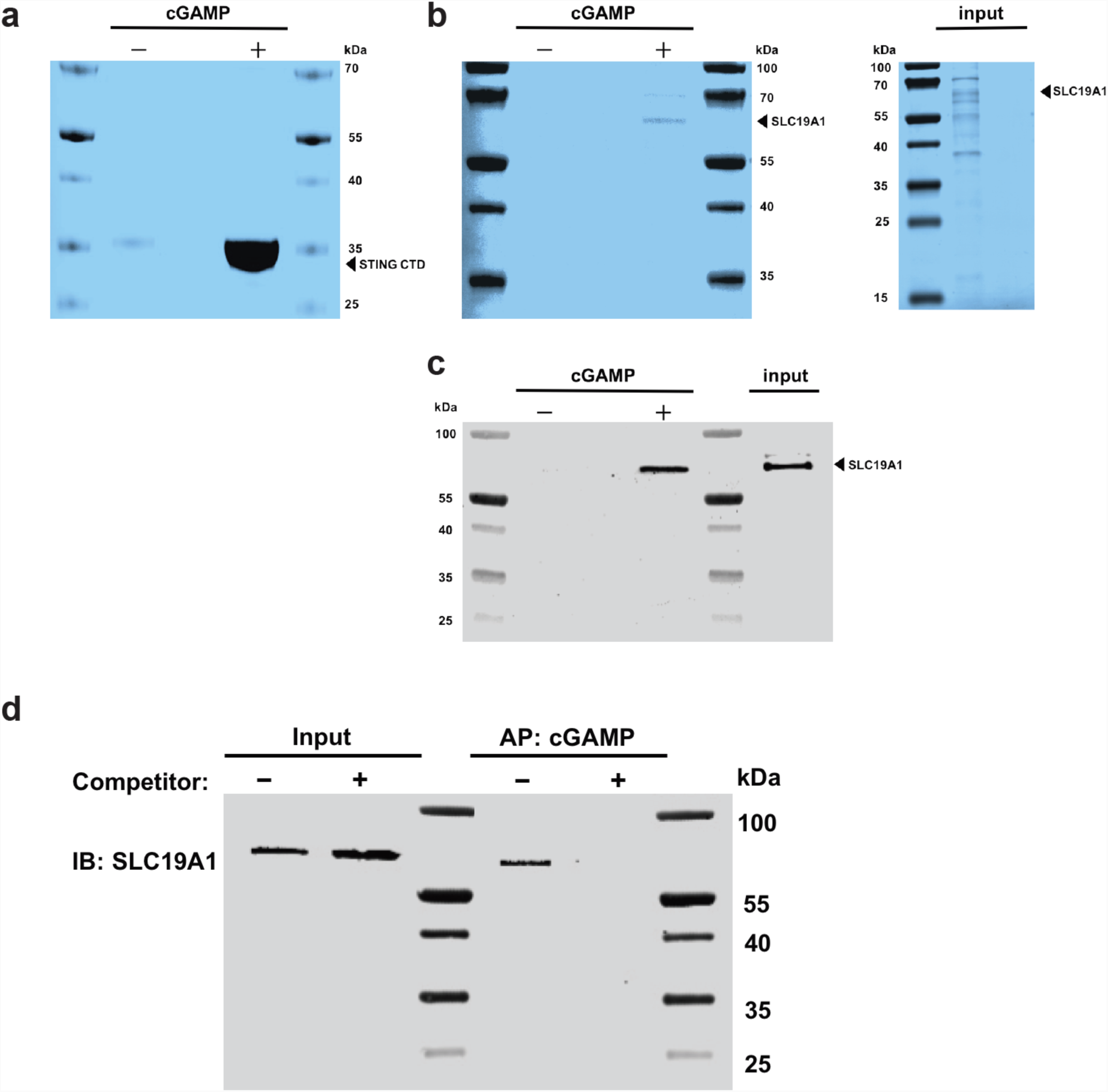
SLC19A1 interacts with 2’3’ cGAMP. **a,** Sodium dodecyl sulfate (SDS)-PAGE analysis followed by Coomassie Blue staining of mSTING-C-Terminal Domain (CTD) pull-downs with 2’3 cGAMP (+) or control (−) Sepharose. **b,** SDS-PAGE analysis followed by Coomassie Blue staining of His-tagged hSLC19A1 pull-downs with 2’3’ cGAMP (+) or control (−) Sepharose as well as the input material following Ni-NTA affinity purification (right panel). **c,** SDS-PAGE analysis followed by Western blot analysis of His-tagged hSLC19A1 pull-downs with 2’3’ cGAMP (+) or control (−) Sepharose as well as the input material following Ni-NTA affinity purification. The two panels were run on the same gel but separated for comparison to the panels in B. **d,** SDS-PAGE analysis followed by Western blot analysis of 8xHis-tagged hSLC19A1 affinity purification (AP) with 2’3’ cGAMP Sepharose in the absence (−) and presence (+) of free, unbound 2’3’ cGAMP (250 µM).

**Figure S9.**
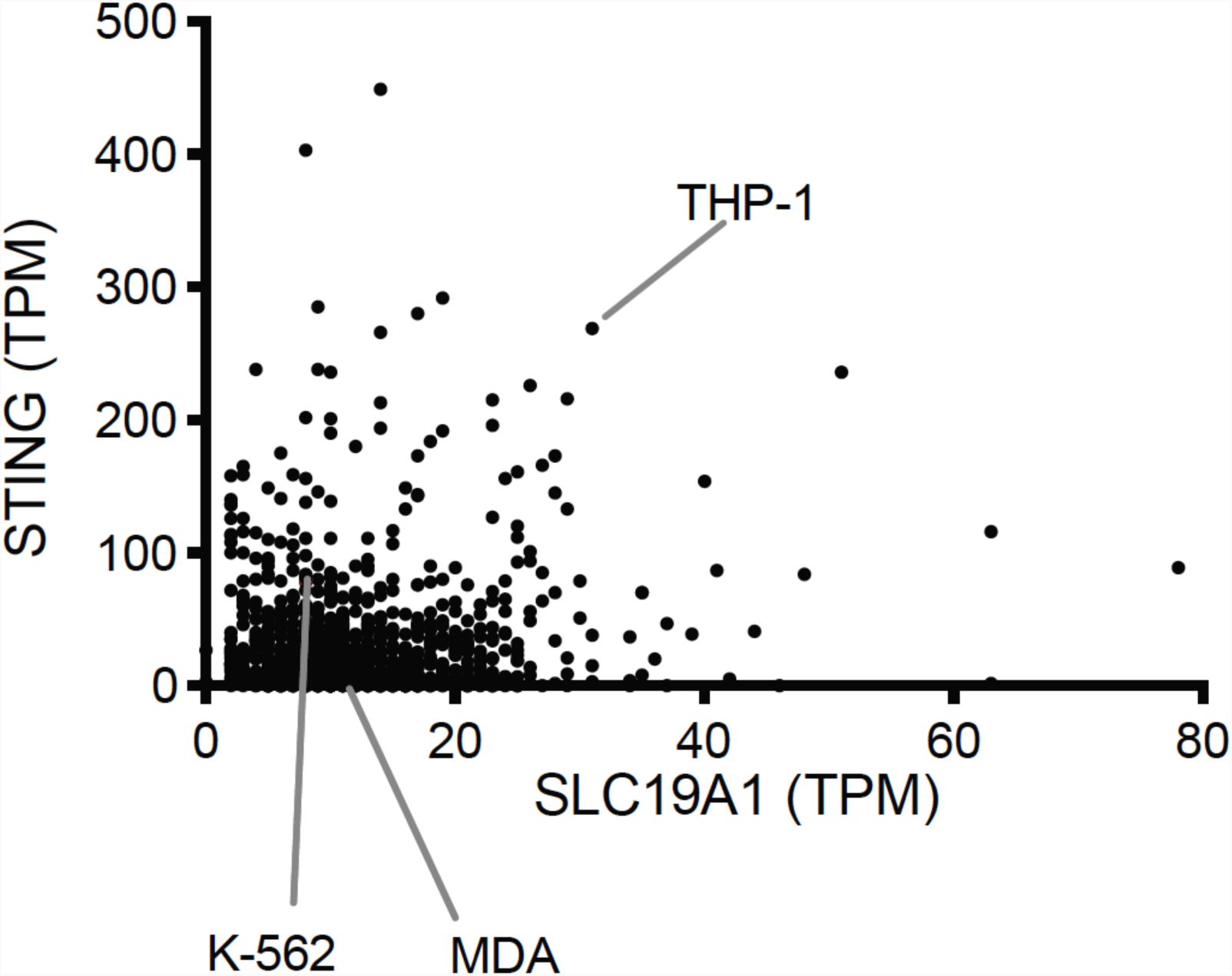
RNA-Seq data of STING and SLC19A1 mRNA expression in 934 human cancer cell lines available at the Cancer Cell Line Encyclopedia. Expression is presented as transcripts per kilobase million (TPM). Data is downloaded from the European Bioinformatics Institute Gene expression Atlas (URL: https://www.ebi.ac.uk/gxa/home). The data set included three of the cell lines we examined, as shown.

**Figure S10.**
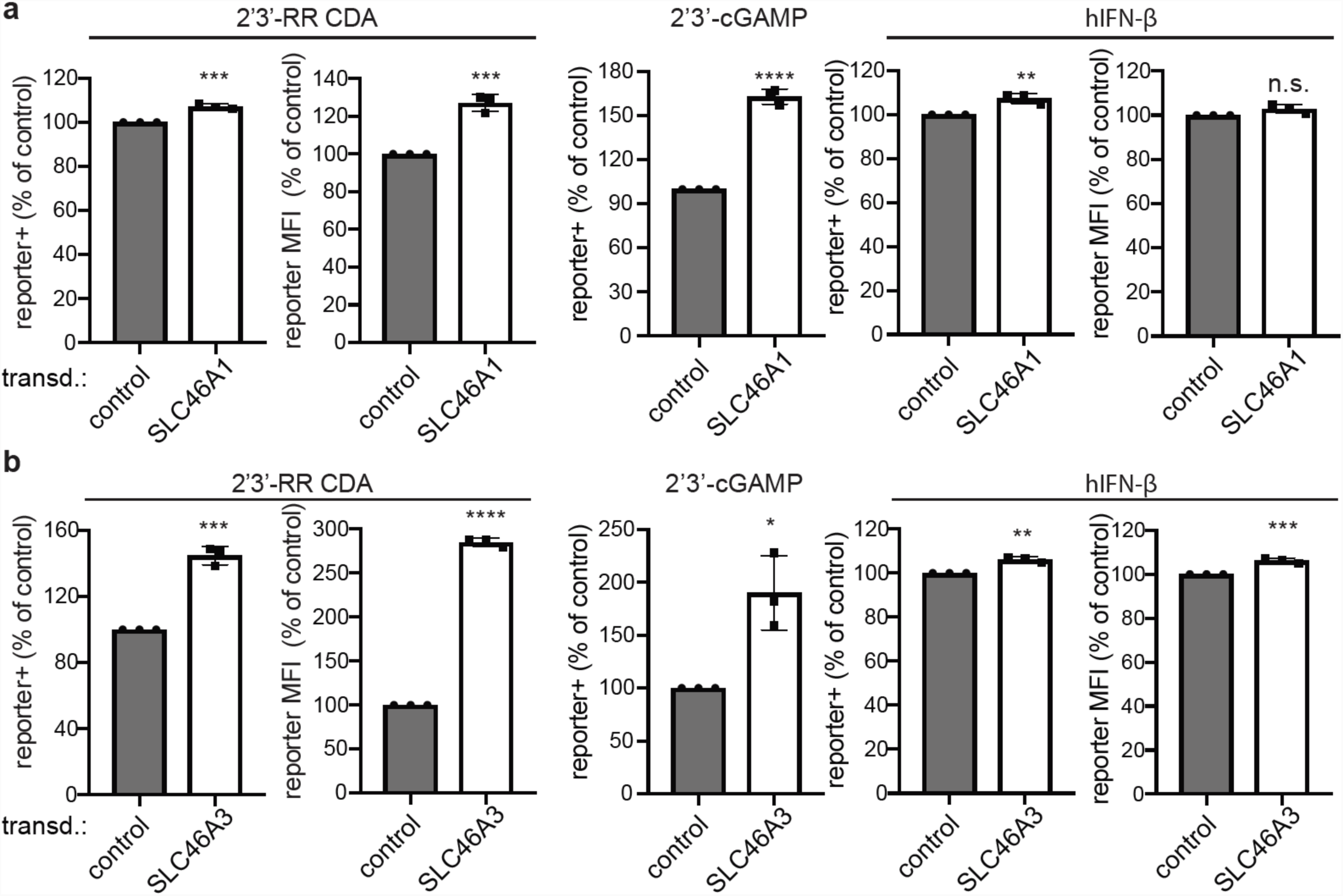
Enforced expression of *SLC46A1* and *SLC46A3* affects the responses of THP-1 cells to CDNs. Control THP-1 cells (transduced with empty expression vector) and *SLC46A1*-transduced THP-1 cells (a) or control THP-1 cells and *SLC46A3*-transduced cells (b) were stimulated with 2’3’-RR CDA (1.25 µg/ml), 2’3’-cGAMP (15 µg/ml) or hIFN-β (100 ng/ml). tdTomato reporter expression was measured by flow cytometry 18-22h after stimulation. Combined data of three independent experiments. Statistical analysis was performed using an unpaired two-tailed Student’s t test. Error bars represent ± SE of independent replicates. \**P* ≤ 0.05; \*\**P* ≤ 0.01;\*\*\**P* ≤0.001; \*\*\*\**P* ≤ 0.0001; n.s. not significant.

**Figure S11.**
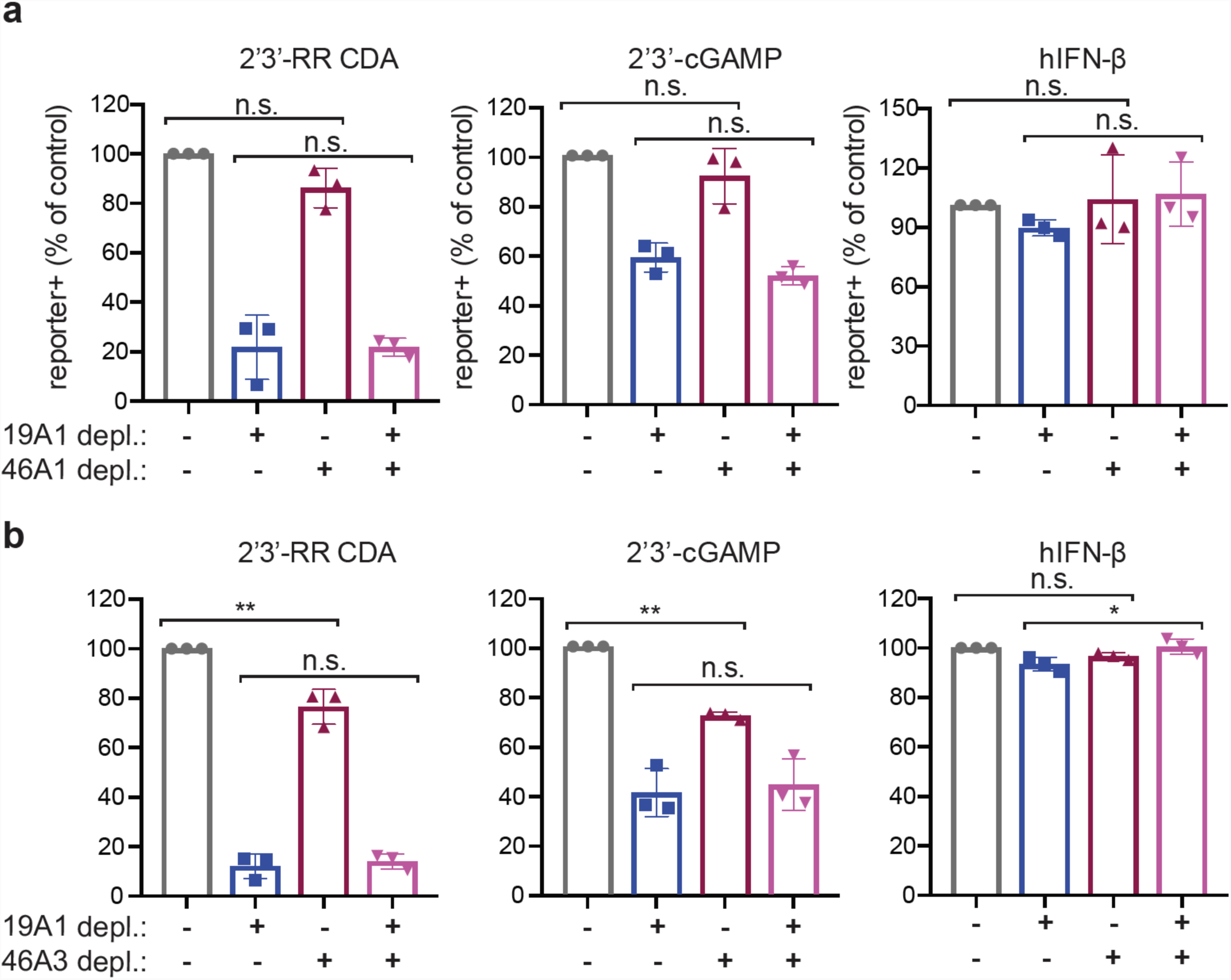
*SLC46A1* or *SLC46A3* depletions, in combination with *SLC19A1* depletion have no additional effect on stimulation by CDNs. THP-1 cells were transduced with non-targeting control CRISPRi gRNAs or *SLC19A1*-targeting CRISPRi gRNA in combination with a second control CRISPRi gRNA or *SLC46A1*-targeting CRISPRi gRNA in (a) or *SLC46A3*-targeting gRNA in (b). Cells were stimulated with 2’3’-RR CDA (1.67 µg/ml), 2’3’-cGAMP (10 µg/ml), or hIFN-β (100 ng/ml). tdTomato reporter expression was measured by flow cytometry 18-22h after stimulation. Combined data of three independent experiments. Statistical analysis was performed using a one-way ANOVA followed by a Tukey’s post-test, comparing only the effects of depleting SLC46A1 (a) or SLC46A3 (b). Error bars represent ± SE of independent replicates. \**P* ≤ 0.05; \*\**P* ≤ 0.01; n.s. not significant.

## References

1. Ishii, K. J. et al. A toll-like receptor-independent antiviral response induced by double-stranded B-form DNA. Nat. Immunol. 7, 40–48 (2006).

2. Stetson, D. B. & Medzhitov, R. Recognition of cytosolic DNA activates an IRF3-dependent innate immune response. Immunity 24, 93–103 (2006).

3. Li, T. & Chen, Z. J. The cGAS–cGAMP–STING pathway connects DNA damage to inflammation, senescence, and cancer. J. Exp. Med. 215, 1287–1299 (2018).

4. Zhang, X. et al. Cyclic GMP-AMP containing mixed Phosphodiester linkages is an endogenous high-affinity ligand for STING. Mol. Cell 51, 226–235 (2013).

5. Sun, L., Wu, J., Du, F., Chen, X. & Chen, Z. J. Cyclic GMP-AMP synthase is a cytosolic DNA sensor that activates the type I interferon pathway. Science 339, 786–91 (2013).

6. Gao, P. et al. Cyclic [G(2′,5′)pA(3′,5′)p] is the metazoan second messenger produced by DNA-activated cyclic GMP-AMP synthase. Cell 153, 1094–1107 (2013).

7. Ablasser, A. et al. CGAS produces a 2′-5′-linked cyclic dinucleotide second messenger that activates STING. Nature 498, 380–384 (2013).

8. Diner, E. J. et al. The Innate Immune DNA Sensor cGAS Produces a Noncanonical Cyclic Dinucleotide that Activates Human STING. Cell Rep. 3, 1355–1361 (2013).

9. Ishikawa, H. & Barber, G. N. STING is an endoplasmic reticulum adaptor that facilitates innate immune signalling. Nature 455, 674–8 (2008).

10. Corrales, L. et al. Direct Activation of STING in the Tumor Microenvironment Leads to Potent and Systemic Tumor Regression and Immunity. Cell Rep. 11, 1018–30 (2015).

11. Marcus, A. et al. Tumor-Derived cGAMP Triggers a STING-Mediated Interferon Response in Non-tumor Cells to Activate the NK Cell Response. Immunity 49, 754–763.e4 (2018).

12. McWhirter, S. M. et al. A host type I interferon response is induced by cytosolic sensing of the bacterial second messenger cyclic-di-GMP. J. Exp. Med. 206, 1899–1911 (2009).

13. Dey, R. J. et al. Inhibition of innate immune cytosolic surveillance by an M.Tuberculosis phosphodiesterase. Nat. Chem. Biol. 13, 210–217 (2017).

14. Woodward, J. J., Lavarone, A. T. & Portnoy, D. A. C-di-AMP secreted by intracellular Listeria monocytogenes activates a host type I interferon response. Science (80-.). 328, 1703–1705 (2010).

15. Barker, J. R. et al. STING-dependent recognition of cyclic di-AMP mediates type I interferon responses during Chlamydia trachomatis infection. MBio 4, 1–11 (2013).

16. Lam, A. R. et al. RAE1 ligands for the NKG2D receptor are regulated by STING-dependent DNA sensor pathways in lymphoma. Cancer Res. 74, 2193–2203 (2014).

17. Ahn, J., Gutman, D., Saijo, S. & Barber, G. N. STING manifests self DNA-dependent inflammatory disease. Proc. Natl. Acad. Sci. U. S. A. 109, 19386–91 (2012).

18. Gao, D. et al. Activation of cyclic GMP-AMP synthase by self-DNA causes autoimmune diseases. Proc. Natl. Acad. Sci. 112, E5699–E5705 (2015).

19. Gall, A. et al. Autoimmunity initiates in nonhematopoietic cells and progresses via lymphocytes in an interferon-dependent autoimmune disease. Immunity 36, 120–31 (2012).

20. Woo, S. R. et al. STING-dependent cytosolic DNA sensing mediates innate immune recognition of immunogenic tumors. Immunity 41, 830–842 (2014).

21. Corrales, L. & Gajewski, T. F. Molecular Pathways: Targeting the Stimulator of Interferon Genes (STING) in the Immunotherapy of Cancer. Clin. Cancer Res. 21, 4774–9 (2015).

22. Corrales, L., McWhirter, S. M., Dubensky, T. W. & Gajewski, T. F. The host STING pathway at the interface of cancer and immunity. J. Clin. Invest. 126, 2404–11 (2016).

23. Sundararaman, S. K. & Barbie, D. A. Tumor cGAMP Awakens the Natural Killers. Immunity 49, 585–587 (2018).

24. Gentili, M. et al. Transmission of innate immune signaling by packaging of cGAMP in viral particles. Science 349, 1232–6 (2015).

25. Bridgeman, A. et al. Viruses transfer the antiviral second messenger cGAMP between cells. Science 349, 1228–32 (2015).

26. Ablasser, A. et al. Cell intrinsic immunity spreads to bystander cells via the intercellular transfer of cGAMP. Nature 503, 530–534 (2013).

27. Xu, S. et al. cGAS-Mediated Innate Immunity Spreads Intercellularly through HIV-1 Env-Induced Membrane Fusion Sites. Cell Host Microbe 20, 443–457 (2016).

28. Chen, Q. et al. Carcinoma-astrocyte gap junctions promote brain metastasis by cGAMP transfer. Nature 533, 493–498 (2016).

29. Hou, Z. & Matherly, L. H. Biology of the major facilitative folate transporters SLC19A1 and SLC46A1. Current Topics in Membranes 73, (Elsevier Inc., 2014).

30. Jansen, G. et al. Sulfasalazine is a potent inhibitor of the reduced folate carrier: Implications for combination therapies with methotrexate in rheumatoid arthritis. Arthritis Rheum. 50, 2130–2139 (2004).

31. Lin, R., Heylbroeck, C., Genin, P., Pitha, P. M. & Hiscott, J. Essential Role of Interferon Regulatory Factor 3 in Direct Activation of RANTES Chemokine Transcription. Mol. Cell. Biol. 19, 959–966 (1999).

32. Brownell, J. et al. Direct, Interferon-Independent Activation of the CXCL10 Promoter by NF-B and Interferon Regulatory Factor 3 during Hepatitis C Virus Infection. J. Virol. 88, 1582–1590 (2014).

33. Donaldson, G. P., Roelofs, K. G., Luo, Y., Sintim, H. O. & Lee, V. T. A rapid assay for affinity and kinetics of molecular interactions with nucleic acids. Nucleic Acids Res. 40, (2012).

34. Zhao, R., Diop-Bove, N., Visentin, M. & Goldman, I. D. Mechanisms of Membrane Transport of Folates into Cells and Across Epithelia. Annual Review of Nutrition 31, (2011).

35. King, K. R. et al. IRF3 and type i interferons fuel a fatal response to myocardial infarction. Nat. Med. 23, 1481–1487 (2017).

36. Ahn, J., Son, S., Oliveira, S. C. & Barber, G. N. STING-Dependent Signaling Underlies IL-10 Controlled Inflammatory Colitis. Cell Rep. 21, 3873–3884 (2017).

37. Plosker, G. L. & Croom, K. F. Sulfasalazine: a review of its use in the management of rheumatoid arthritis. Drugs 65, 1825–49 (2005).

38. Kozuch, P. L. & Hanauer, S. B. Treatment of inflammatory bowel disease: A review of medical therapy. World J. Gastroenterol. 14, 354–377 (2008).

39. Wahl, C., Liptay, S., Adler, G. & Schmid, R. M. Sulfasalazine: A potent and specific inhibitor of nuclear factor kappa B. J. Clin. Invest. 101, 1163–1174 (1998).

40. van de Weijer, M. L. et al. A high-coverage shRNA screen identifies TMEM129 as an E3 ligase involved in ER-associated protein degradation. Nat. Commun. 5, 3832 (2014).

41. van Diemen, F. R. et al. CRISPR/Cas9-Mediated Genome Editing of Herpesviruses Limits Productive and Latent Infections. PLoS Pathog. 12, e1005701 (2016).

42. Horlbeck, M. A. et al. Compact and highly active next-generation libraries for CRISPR-mediated gene repression and activation. Elife 5, 1–20 (2016).

43. Kampmann, M., Bassik, M. C. & Weissman, J. S. Functional genomics platform for pooled screening and generation of mammalian genetic interaction maps. Nat. Protoc. 9, 1825–47 (2014).

44. Gilbert, L. A. et al. Genome-Scale CRISPR-Mediated Control of Gene Repression and Activation. Cell 159, 647–661 (2014).

45. Kampmann, M., Bassik, M. C. & Weissman, J. S. Integrated platform for genome-wide screening and construction of high-density genetic interaction maps in mammalian cells. Proc. Natl. Acad. Sci. 110, E2317–E2326 (2013).

46. Sadlish, H., Williams, F. M. R. & Flintoff, W. F. Functional Role of Arginine 373 in Substrate Translocation by the Reduced Folate Carrier. J. Biol. Chem. 277, 42105–42112 (2002).

47. Huynh, T. N. et al. An HD-domain phosphodiesterase mediates cooperative hydrolysis of c-di-AMP to affect bacterial growth and virulence. Proc. Natl. Acad. Sci. U. S. A. 112, E747–56 (2015).

48. Sureka, K. et al. The cyclic dinucleotide c-di-AMP is an allosteric regulator of metabolic enzyme function. Cell 158, 1389–1401 (2014).

49. McFarland, A. P. et al. Sensing of Bacterial Cyclic Dinucleotides by the Oxidoreductase RECON Promotes NF-κB Activation and Shapes a Proinflammatory Antibacterial State. Immunity 46, 433–445 (2017).

50. Ray, A. & Dittel, B. N. Isolation of mouse peritoneal cavity cells. J. Vis. Exp. (2010). doi:10.3791/1488

